# Topographical and spectral signatures of infant and adult movement artifacts in naturalistic EEG

**DOI:** 10.1101/206029

**Authors:** Stanimira Georgieva, Suzannah Lester, Meryem Nazli Yilmaz, Sam Wass, Victoria Leongi

**Affiliations:** Department of Psychology, University of Cambridge; University of East London; Nanyang Technological University, Singapore

**Keywords:** EEG, Signal distortion, Motion artifacts, Infants

## Abstract

Electroencephalography (EEG) is perhaps the most widely used brain-imaging technique for paediatric populations. However, EEG signals are prone to distortion by motion. Compared to adults, infants’ motion is both more frequent and less stereotypical yet motion effects on the infant EEG signal are largely undocumented. Here, we present a systematic assessment of naturalistic motion effects on the infant and adult EEG signal. In Study 1, we documented the prevalence of 27 naturally occurring facial and body motions by video-coding five mother-infant pairs during naturalistic play. In Study 2, we elicited a subset of the most common facial, limb and postural motions from one adult and one infant actor while their EEG was recorded. In Study 3, we recorded EEG signals from a larger group of 12 infants whilst they produced the same motions spontaneously. Our findings from Study 2 suggested that adult movements mainly generated *increases* in spectral power relative to resting state, primarily at peripheral sites and in delta and high-beta frequency bands. In infants, both elicited motions (N=1, Study 2) and spontaneously-occurring motions (N=12, Study 3) produced *decreases* in theta, alpha and beta power over central regions and *increased* beta/delta power at peripheral sites. We also observed that infants’ jaw and upper limb movements generated more pronounced EEG artifacts than lower limb movements. It is intended that this work will inform future development of methods for addressing EEG motion-related artifacts and support wider use of naturalistic paradigms in social and developmental neuroscience.

## 1 INTRODUCTION

### 1.1 Motion in EEG measurements

Electroencephalography (EEG) is a widely used brain imaging technique for both adult and paediatric populations, owing to its low risk to the individual (Teplan, 2002) and ease of application (De Haan, 2013). In particular, rising interest in the neural processes that play a critical part in early emotional, social, and cognitive development has led to an increased use of EEG with infants and young children (Saby and Marshall, 2012). For decades, research in infants and young children has employed EEG as a methodology to understand the neural processes involved in numerous aspects of early cognitive development (Maguire et al., 2014). Examples include early language acquisition (Kuhl, 2010), visual attention and object recognition in infants (Reynolds, 2015), attention (Richards et al., 2010), face and emotion processing (Batty et al., 2011) and auditory processing (Telkemeyer et al., 2011).

However, EEG recordings are highly prone to distortion by both biological factors (such as electromyogenic activity) and non-biological factors (such as electrical line noise) (Nathan and Contreras-Vidal, 2015). In particular, artifacts induced by motion (such as head motion, jaw motion, or blinking) are a major and common source of EEG distortion. These distortions can result in misinterpretation of underlying neural processes or sources; or to the inaccurate detection and diagnosis of brain disorders (Guerrero-Mosquera et al., 2012). For adult populations, motion artifacts can be avoided or minimised by direct instruction, such as asking participants to only swallow and blink during defined periods and asking them to avoid significant head and facial muscle contractions during recordings (Reis et al., 2014). While direct instruction may be efficacious for reducing movements in healthy adult participants, this strategy is less effective for clinical and paediatric populations whose ability to understand and comply with verbal instruction is greatly reduced. Young infants present a particular challenge in this regard, as they have a high natural tendency for movement, which cannot be constrained by instruction. Indeed, it is widely acknowledged that EEG recordings produced by infants and young children are heavily contaminated by various motion artifacts, including gross motor movements and eye blinks (Bell and Cuevas, 2012). Motion artifacts can be a major confound to the interpretation of neural data, particularly if the type or quantity of motion differs systematically across the experimental conditions or groups of interest.

One common strategy used in infancy paradigms is to reduce motion indirectly through attentional capture – that is, the experimenter monitors the infants’ attentional state through a video feed and only delivers experimental stimuli during attentive periods when the infant is relatively still. However, as exemplified in Table S1 (Supplementary Materials), our own studies suggest that even when contingently delivered screen-based stimuli are used (including cartoons, real language and artificial language stimuli), infant movement (i.e. facial, limb or postural movement) is still present throughout 60-70% of the total stimulus presentation time. In more naturalistic paradigms, in which infants are not watching a computer monitor but engaged in social interaction, we might expect that artifacts will be more prominent still.

### 1.2 Naturalistic social paradigms

The necessity for ecological validity in experimental developmental psychology has been emphasised for decades (Tunnell, 1977; Fabes et al., 2000). It is accepted that the combination of experimental and naturalistic research methods offers more complete insight into child development (Dahl, 2016). Observational assessments of behaviour in the ‘real world’, such as in the home or school environment, have higher ecological validity than assessments that occur within structured experiments in laboratory settings. However, real-world measurements also carry the disadvantage of less (or no) control over key environmental variables that can affect behaviour, leading to increased inter-subject variability. As a compromise, naturalistic laboratory settings allow for some controlled task-based variation between participants, and facilitate the emergence of more natural behaviour whilst at the same time minimising environmental variation (Tamis-LeMonda et al., 2017; Noris et al., 2012). The use of naturalistic methods is more common in social science research than in the neurosciences. For example, in education research, naturalistic methods are recognised as providing a solution to the problems of diminished construct validity and poor generalisability associated with laboratory settings (Smith, 1982). In neuroscience research, by contrast, the necessity for carefully controlled experiments has historically outweighed the importance of ecological validity. However, across a number of neuroscience sub-fields, such as developmental and social neuroscience, the balance is beginning to shift in favour of paradigms with greater ecological validity.

For example, Dumas et al. (2014) argue that many social coordination phenomena cannot be observed in the lab without the presence of at least two individuals, and their natural social interaction patterns. In this, and other hyperscanning studies, behavioural and neural (e.g. EEG) data is typically simultaneously acquired from two or more individuals who are engaged in naturalistic social interaction, such as gesture imitation or gaze-following (Dumas et al., 2010, 2011). The resulting neural coupling between brains (e.g. centro-parietal alpha synchrony) has been found to reflect continuous mutual adaptations of behaviour in participant dyads (Dumas et al., 2010). In another example of EEG use during naturalistic social interactions, Lindenberger et al. (2009) demonstrated that pairs of guitarists who play together also show cross-brain theta phase alignment, which is modulated by the lead guitarist’s behavioural gestures as well as the onset of play itself. Finally, Babiloni et al. (2007) used high-density EEG on participants engaged in the “Prisoner’s Dilemma” – a game that requires cooperation and competition between the players, therefore engaging higher cognitive functions such as memory and decision-making. Based on participants’ (naturalistic) social interaction patterns, the authors were able to identify distinct neural functional connectivity patterns associated with social cooperation as opposed to conflict or competition. In all of these neuroimaging studies, a tension exists between the ecological benefits conferred by naturalistic social interaction, and the generation of EEG artifacts from participants’ social behaviour (e.g. facial and gesticulatory movements).

### 1.3 Examples of common EEG motion artifacts

Neural activity at the scalp level is low in amplitude compared to other sources of electrical activity, such as electromyogenic potentials and environmental electrical noise (i.e. a low signal-to-noise ratio). Non-biological artifacts (such as line noise) often show a different frequency signature from that of neural signals and can therefore be relatively easily isolated by filtering techniques. When adequate grounding is used in an electrical interference-free environment, the major source of concern are biological artifacts such as muscle contractions, whose frequency content overlaps significantly with that of neural signals (and therefore cannot be easily removed by filtering). Amplification that is applied to the neural signal also amplifies non-neural contaminants, and therefore does not improve neural signal detection. There is already some understanding of the effects of eye and jaw motions (as described below). However, this literature pertains almost exclusively to adults, and virtually nothing is known about the nature of motion artifacts in *infant* EEG signals.

One common source of movement-related artifacts in adult EEG are those produced by the eyes. A single eye movement can produce a number of artifacts that arise from different mechanisms (e.g. eye rotations and blinks) and differ in their amplitude and spectral properties (Plöchl et al., 2012). Further, eye movements can introduce systematic biases in both adult (Yuval-Greenberg et al., 2008; Keren et al., 2010) and infant (Koster, 2016) EEG analyses. For example, in a concurrent EEG and eye-tracking paradigm, Yuval-Greenberg et al. (2008) showed that the most likely source of the induced gamma-band EEG response (iGBR) – a neural component associated with visual object representation, recognition and attention – was ocular rather than neural. They noted that even at the single trial level, activity that was spatially and temporally consistent with the iGBR only occurred on saccade trials and was time-locked to the saccade. Koster (2016) highlighted that a similar problem may exist for infant EEG analyses utilising activity in the 25-35 Hz gamma range. However, it has not yet been clarified whether infant microsaccades also generate similar EEG effects.

Jaw movements are another major source of EEG artifacts. In experimental paradigms, jaw motion commonly occurs as a corollary of speech production (Ganushchak et al., 2011). Jaw motion causes significant distortion to EEG signals due to facial myogenic potentials originating from contractions of the frontalis and temporalis muscles when tensing or clenching the jaws (Sweeney et al., 2012). Speech-related articulatory motions are known to reduce the signal-to-noise ratio of neural signals that relate to cognition (Brooker & Donald, 1980). For instance, the myogenic potential generated by the temporalis muscle, used for closing the lower jaw, spreads widely over the scalp locations that correspond to the frontal/temporal/parietal junction of the brain, generating large broadband artifacts in the EEG signals measured over these regions (Brooker & Donald, 1980).

Other myogenic EEG artifacts can arise from involuntary movements that support the physiological functioning of the body, such as heartbeat and respiratory torso movements. Such involuntary movement can be monitored using a devoted channel, such as an electrocardiogram (ECG), which can significantly improve automated detection and removal strategies (Klados et al., 2011). By contrast, voluntary movement, such as motion generated by the jaw, head, body and limb, and less frequently addressed in the literature. For example, in an extensive review of methods for EEG artifact detection and rejection, Islam et al. (2016) found that across 46 publications that were reviewed, over 70% focussed solely on automatic movements, with the rest only partially addressing forms of voluntary action. However, this lack of knowledge about the effects of voluntary motion presents a real problem for naturalistic developmental research, as voluntary motion in these situations is both prevalent and unavoidable.

### 1.4 Current strategies for addressing motion artifacts

There are two major approaches to addressing the problem of movement-related artifacts in EEG data. Researchers typically (1) exclude artifact-contaminated segments by employing strict rejection procedures/thresholds; or (2) attempt to remove artifacts from data using correction procedures such as independent component analysis (ICA) (Gwin et al., 2010). The first approach (artifact exclusion) is conservative but may entail considerable data loss (see for example, Table S1). Therefore, there is increasing interest in correction procedures that permit the accurate identification and removal of artifacts from EEG data without significant compromise to the integrity of underlying neural activity.

Several methods have been proposed for the detection and removal of physiologically generated artifacts from the EEG signal. These include the use of linear regression (Klados et al., 2011), and blind source separation (BSS) based on Canonical Correlation Analysis (CCA, De Vos et al., 2010) or Independent Component Analysis (ICA, Porcaro et al., 2015) to separate cortical sources from electromyographic (EMG) responses. However, none of these methods is able to completely remove motion artifacts from the EEG signal and may even remove some genuine neural activity of interest. As noted by Islam et al. (2016), current artifact detection-removal methods are sub-optimal because these methods typically only address a single artifact class and necessitate dedicated reference channels, and moreover, frequently result in overcorrection. For example, a common approach to addressing some classes of stereotypical artifacts (such as eye blinks) is to include an observed reference channel that independently measures the artifact signal (ie. EOG for eye muscles, ECG for heartbeat). Next, linear regression or ICA may be employed to estimate the similarities between the EEG signal and the reference signal, permitting removal of the artifact estimate from the EEG signal (Klados et al., 2011). While this approach can be successful for highly-stereotypical artifacts (such as heartbeats), it fails for less stereotypical artifacts, such as EMG responses. Further, the placement of additional reference channels (e.g. under the eyes) may not be well tolerated by paediatric and clinical subjects.

Unlike regression, ICA does not rely on the presence of a reference signal (although this can improve performance, see Plöchl et al., 2012), and is therefore a widely used approach for artifact removal (Chaumon et al., 2015). ICA is a BSS-based technique which additionally assumes that all sources generating the measured signal are orthogonal to each other (Islam et al., 2016). BSS techniques, in general, aim to estimate the individual contributors to a mixed signal as a linear summation between all sources, with the addition of noise. ICA can be useful for isolating brain activity from EMG artifacts (McMenamin et al., 2010). However, Shackman et al. (2009) caution that the efficacy of ICA in artifact removal is highly contingent on the user’s own expertise (e.g. understanding which components, and the optimal number of components to remove).

Other less frequently used methods for artifact removal are Empirical Mode Decomposition (EMD), Adaptive filtering (AF), and Principal Component Analysis (PCA), (see Islam et al., 2016 for a review). EMD decomposes a signal into a sum of band-limited components with well-defined instantaneous frequencies (Molla et al., 2012). EMD does not assume linearity or stationarity of the signal and is therefore well suited for EEG. However, the method is computationally demanding and has not yet been tested in empirical studies, therefore little can be concluded about its efficacy (Islam et al., 2016). In adaptive filtering, a linear transfer function filter is used, whose weights are defined by an adaptive optimization algorithm (Widrow & Stearns, 1985). The weights are adapted to filter out the artifactual source based on a reference channel, which is assumed to be independent from the EEG signal. This method has been successfully applied to EOG artifact removal (Islam et al., 2016). PCA is a spatial filtering procedure in which the EEG signal is orthogonally transformed into linearly uncorrelated components (Cohen, 2014). However, as the resulting PCA components are ordered by the amount of variance explained, difficulties arise when neural and myogenic signals have similar amplitudes. Therefore, the efficacy of PCA as compared to other decomposition techniques has been questioned (Turnip & Junaidi, 2014).

Newer machine learning approaches have also been employed for automatic classification of artifactual and non-artifactual segments of EEG signals. For example, O’Regan et al. (2010) trained a classifier to segregate artifactual and neural EEG signals using a database of over 300 minutes of head motion contaminated EEG data. In their study, 19 healthy participants were instructed to perform 32 types of head actions that had previously been related to distortions in ambulatory EEG, including head shaking, rolling, nodding, jaw clenching, lowering and raising of eye-brows. The EEG signal was epoched into 0.25, 0.5, 0.75, 1.0, 1.5, 2, 3 and 4-second windows, and features were selected based on measures of Mutual Information. Next, a classifier using linear discriminant analysis was trained from a random selection of 20% of the data. Results indicated that the selected feature set was sufficient for the classifier to be able to distinguish between head motion-contaminated and clean EEG with 66% accuracy, and the performance of the classifier was highest for 1.0s and 1.5s window lengths (75.66 % and 76.49%, respectively).

Similarly, Lawhern et al. (2012) investigated methods for automatic detection and classification of EEG artifacts generated by different types of jaw and eye movements. Seven participants provided 20 repetitions of each movement in time to a pacing tone (one tone every two seconds). A baseline (no artifact) condition was also recorded. Epochs of 500ms length were extracted to allow for variability in the duration of movements, ensuring that the full time-course of each artifact was present in each epoch. An autoregressive model using a Maximum Likelihood Estimator was used to estimate features for a support vector machine classifier. The procedure was reportedly successful in differentiating between broad classes of artifacts such as those generated by jaw and eye movements; but it tended to group together artifacts from a similar or common source, such as jaw clenching and chewing movements. However, the error rate of falsely classifying epochs with no artifacts was low (the reported average accuracy for 5 out of 7 participants was over 96%, and over 81% for the remaining 2). Therefore, newer machine learning approaches may, in future, have strong utility for the detection and removal of more complex classes of motion artifacts (such as muscular artifacts from head or jaw motion), where more traditional methods have been less successful. It is anticipated that the data from the current study could be used, in future, to inform the development of such new tools for artifact removal.

### 1.5 Study aims

As the removal of artifacts from EEG data is restricted by current methodological limitations, and clinical and infant populations are unable to comply with directions to reduce movement to lessen distortion of the signal, there is a clear need for research to report how these artifacts distort EEG data. Accordingly, the major aim of this study is to investigate the topological and spectral features of motion-related EEG artifacts in adults and infants, as compared to resting state EEG measurements. We set out to create a detailed description of typical motion-related artifacts, and to identify the spectral signatures and topographies of these artifacts in the major EEG frequency bands of interest (0.05 - 20 Hz).

Three studies were conducted. Study 1 was a behavioural study that aimed to identify the most prevalent motions in a naturalistic play setting where adult (mother) and infant participants interacted with toys in a social or non-social context. We identified the common motions produced during naturalistic play and investigated how these differed between social and non-social experimental conditions. Study 2 was an EEG study in which selected movements from Study 1 were intentionally modelled by/elicited from one adult and one infant actor. We examined the effects of the elicited facial, limb and postural movements on the adult’s and infant’s recorded EEG signals, relative to resting state. Finally, Study 3 measured the EEG signal of a larger group of infants whilst they produced the same movements spontaneously. All studies were approved by the Cambridge Psychology Research Ethics Committee, and parents provided written informed consent on behalf of their children.

## 2 STUDY 1

### 2.1 Methods

#### 2.1.1 Participants

Five infants (1M, 4F) participated in Study 1 along with their mothers. Infants were aged 374.80 days on average (SD = 74.42 days). Four of the five mothers were native English speakers, and one was native Dutch speaker but her infant was exposed daily to English. The mean age for adults was 33.00 years (SD = 3.03 years). All adult participants reported no neurological problems and normal hearing and vision for themselves and their infants.

#### 2.1.2 Materials

The set of toys were appropriate for the infants’ age and included toys of differing shapes, textures and colours to encourage infants’ interest in playing with them.

#### 2.1.3 Protocol

The purpose of this study was to describe and compare the movements produced by the adult and the infant while they were playing with objects separately (separate play, hereafter SP) or jointly (joint play, hereafter JP). In the experimental setup, the infant sat in a high chair, with the adult facing him/her across a table. Joint play and solo play sessions each lasted approximately ten minutes and were conducted in a counterbalanced order across participants. During both conditions, an experimenter ensured that participants were playing as instructed, and she provided new toys simultaneously to both adult and infant as required to sustain their attention and interest. The experimenter explicitly avoided making prolonged social contact with either participant.

##### Solo play (SP) condition

A 40 cm high screen was set up across the table, so that infant could see the adult, but neither participant had full visibility of each other’s face (this was necessary to ensure that play activity was functionally separate, but the infant did not become distressed). Figure 1a shows an example of the experimental set-up. Each participant was given a toy to play with independently, and adult participants were asked to direct their full attention to the toy whilst ignoring the infant. Adults and infants were given the same toy to ensure that their visual and tactile experiences would be similar.

**Figure 1.**
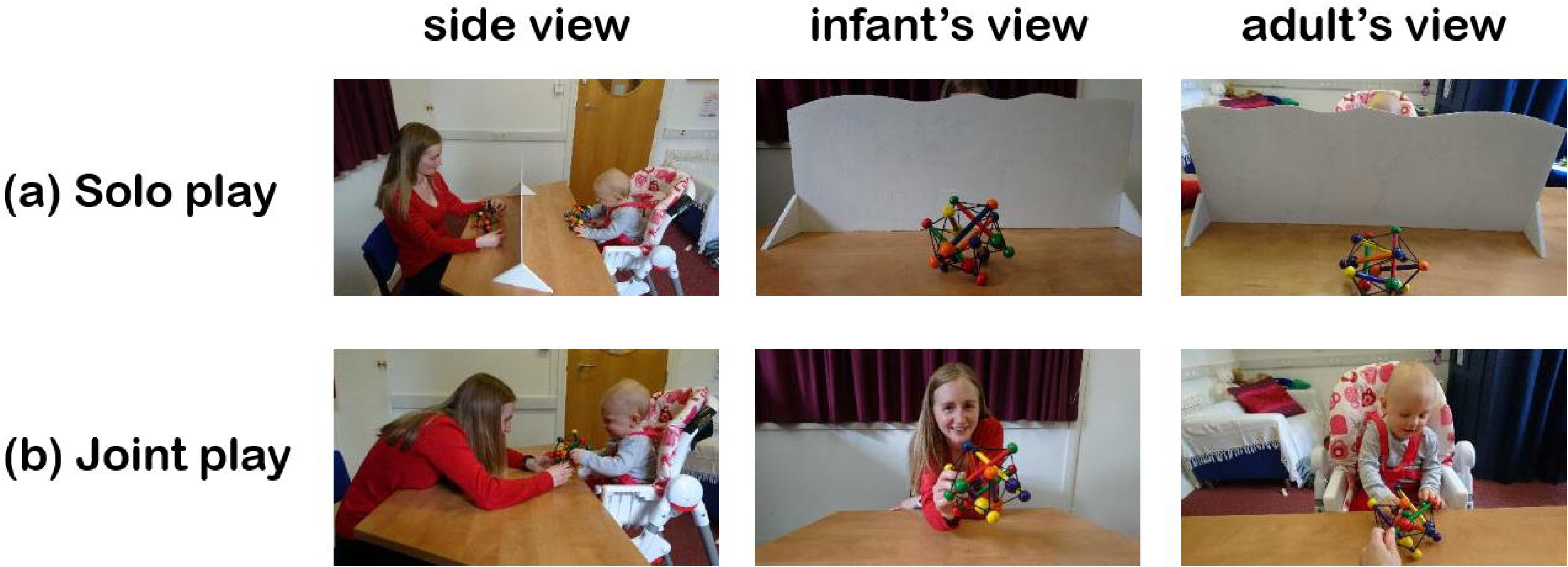
Example of the (a) solo play and (b) joint play experimental set-up showing the (left column) side view (middle column) infant’s view; and (right column) adult’s view.

##### Joint play (JP) condition

For the JP condition, adult participants were asked to play with their infant using the same toys provided in the SP condition (see Figure 1b). In this condition, the adult was requested to actively engage the infant’s attention using the toy objects during joint play. However, the adult was asked to play *silently* with her infant for two reasons. First, we were primarily interested in facial and body movement-related artifacts (directly related to naturalistic play and social interaction) rather than speaking-related artifacts *per se*. Second, during the solo play condition, adults were not speaking.

In both conditions, therefore, the adults were not speaking. However, infants occasionally made vocalisations in both conditions, which were included in our analysis.

#### 2.1.4 Video recordings

To record the actions of the participants, two Logitech High Definition Professional Web-cameras (30 frames per second) were used, directed at the adult and infant respectively. Afterwards, each video recording was manually coded for the behaviours of interest.

#### 2.1.5 Motion coding

Videos were first reviewed to qualitatively identify different classes of facial and body motions that were produced by the adult and the infant and could potentially introduce artifacts into the EEG signal. This analysis was used to devise a coding scheme encompassing 27 different facial and body motions as summarised in Table I below and detailed further in the SM (Sections 2.1 - 2.3). All infants and adults were scored on all of the items on the coding scheme although some of them were more characteristic of the infants, while others of the adults.

Each participant’s video was parsed into 5s-long non-overlapping epochs. Within each epoch, the occurrence of each motion was coded as being either present [code = 1] or absent [code = 0]. The total frequency of occurrence for each motion over the entire session was then computed for each participant. Co-occurrences of different motions (e.g. chewing and small hand movement) were frequently observed. From the coded frequency data, comparative statistics were computed to identify (1) commonly-occurring motions in each condition, and (2) motions that differed between social and non-social conditions. An example of the coding arising from a single participant pair is given in Figure S1 of the Supplementary Materials.

### 2.2 Results

As shown in Figure 2, at least one type of motion was present in over 95% of the time epochs during all tasks. A more detailed breakdown of the occurrence of each class of motions is provided in Tables S1-S3 of the SM. We also noted that the motions overlapped with each other in time (e.g. Figure S1), and infrequently was any one motion observed on its own. To assess whether the total prevalence of motion differed between participants or conditions, a 2 (Participant: Adult, Infant) × 2 (Play Condition: Solo, Joint) ANOVA was conducted. The results showed no significant main effect for Participant, F(1,16) = 3.31, p=.09, or for Play Condition, F(1,16) = 2.92, p=.11. The interaction between Participant and Play Condition was also non-significant, F (1,16) = 3.01, p=.10. Therefore, we observed equal proportions of motion contamination across both play conditions, and for infants’ and adults’ artifacts, when all motion types were pooled together.

**Figure 2.**
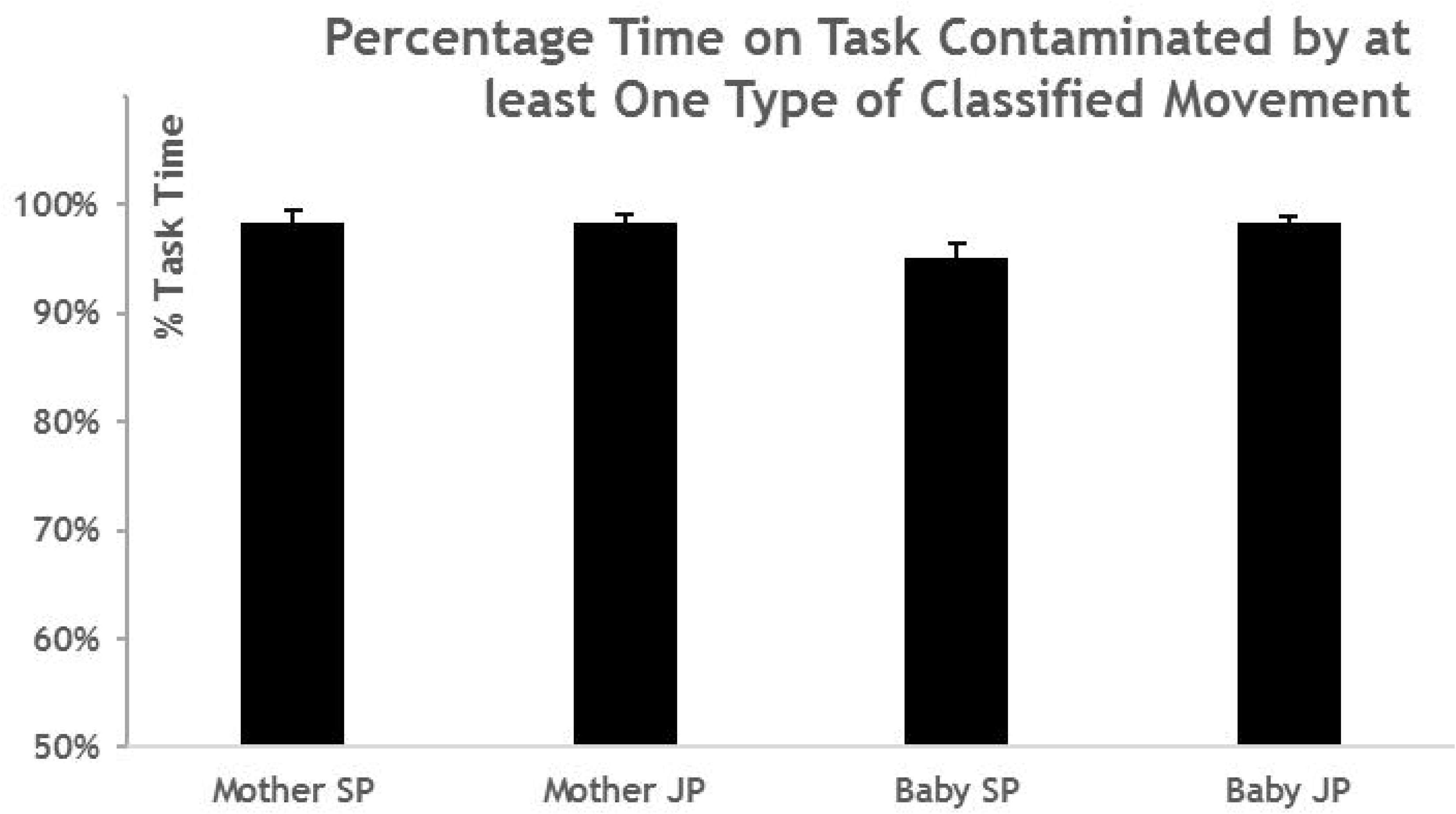
Proportion of the time during Study 1 when at least one movement was observed. Error bars show the standard error of the mean.

Comparative statistics were then computed to identify the most commonly occurring motions in each condition; and to assess whether the prevalence of these most common motions that differed between social (Joint Play, JP) and non-social (Solo Play, SP) conditions. Commonly occurring motions were defined as motions that occurred at least 10% or more of the time (regardless of whether they occurred individually, or in conjunction with other motions). For adult participants, the most common motions were: Up-Down Neck, Lip, Jaw, Small Hand, Large Arm, Leaning Forward and Leaning Back movements (Figure 3). From inspection of Figure 3, the prevalence of all these motions indeed differed between social and non-social conditions, apart from Up-Down Neck Movements. In general, motion was more frequently observed during Joint Play than Solo Play, with the exception of small hand movements that were more frequent during Solo Play. This pattern was confirmed by a series of paired t-tests (p-values uncorrected); Lip Movement: t(4) = 2.70, p =.054; Up and Down Neck Movements: t(4) = 1.92, p=.13; Jaw Movements: t(4) = 3.10, p<.05; Small Hand Movements: t(4) = 4.19, p<.05; Large Arm Movement: t(4) = 5.59, p<.01; Leaning Forward: t(4) = 3.40, p<.05; Leaning Back: t(4) = 4.17, p<.05. However, after applying a (conservative) Bonferroni-correction of p-values (alpha = .05), only Large Arm Movements still showed a significant increase in prevalence for the Joint Play condition.

**Figure 3.**
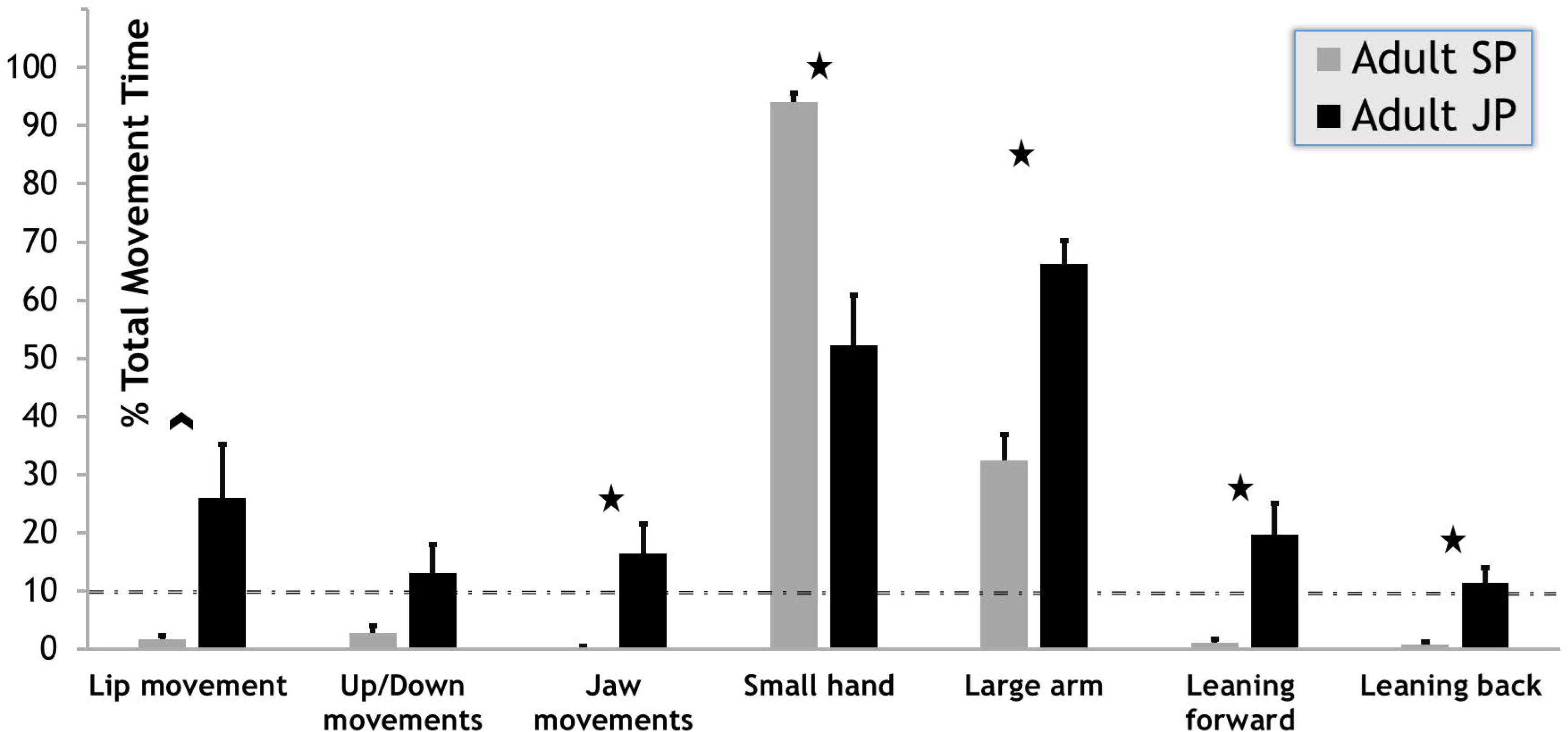
Prevalence of adult motions occurring over 10% of the time in each experimental condition. *p<.05, ^p=.054 (uncorrected)

For infants, the most common movements included Talking, Chewing, Whining, Side-to-Side neck movements, Up and Down neck movements, Small Hand, Small Foot, Large Arm and Large Leg movements (Figure 4). Again, the frequency of each one of these movements was compared between the two play conditions using paired t-tests. Only Chewing, and Up and Down Neck Movements differed significantly in prevalence between Solo and Joint Play conditions, t(4) = 3.68, p<.05 and t(4) = 2.81, p<.05, respectively (pvalues uncorrected, no significant difference for either after Bonferroni correction). There was also a marginal difference between conditions for Side to Side Movements (t(4) = 2.76, p=.051). Unlike the adult, more of these infant motions occurred during Solo Play than during Joint Play. The remaining motions showed no statistical difference in their frequency of occurrence between the two play conditions: talking, whining, small hand movements, large arm movements, small foot movements, large leg movements (all p>.25).

**Figure 4.**
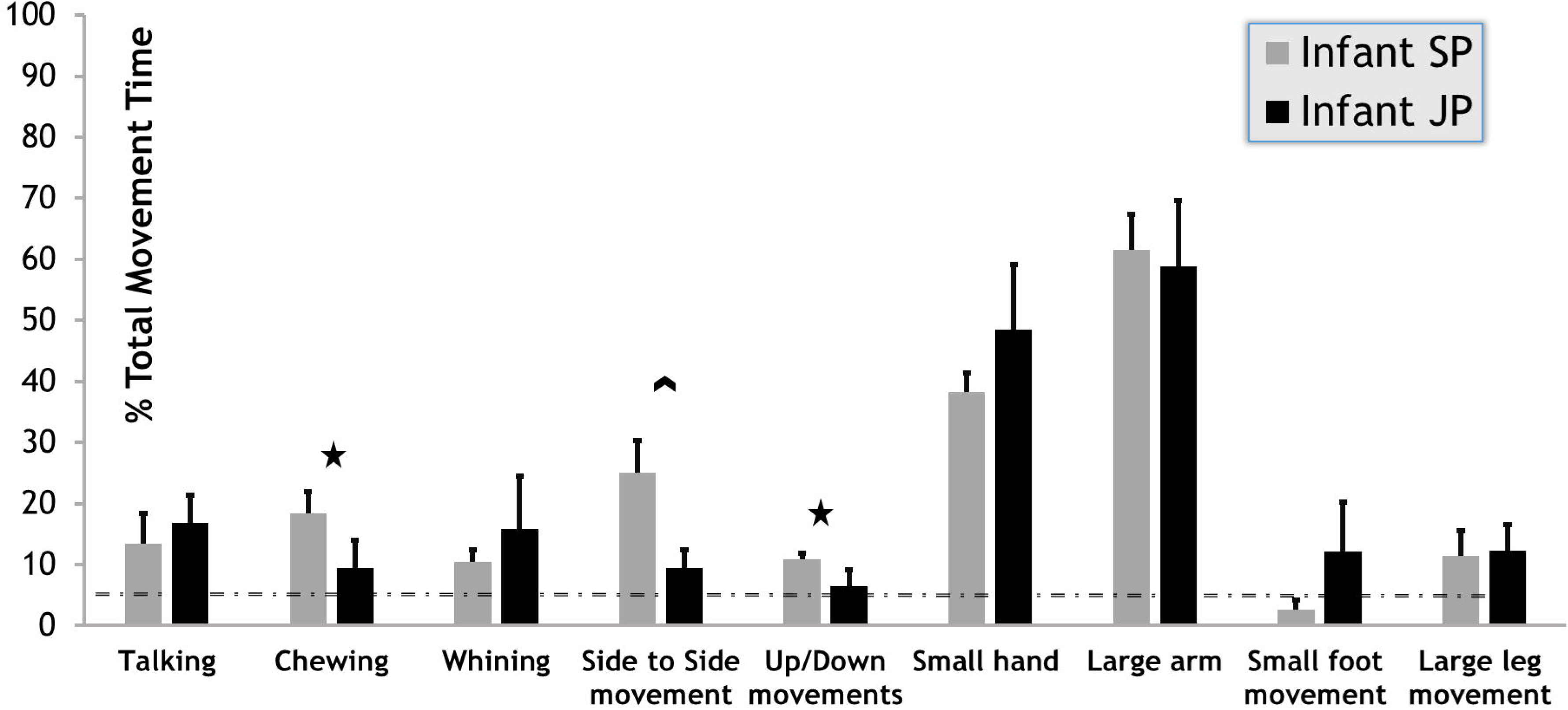
Prevalence of infant motions occurring over 10% of the time in each experimental condition. *p<.05; ^p=.051 (uncorrected)

### 2.3 Study 1 discussion

The aim of Study 1 was to identify the most prevalent types of movements that Adult and Infant participants produced whilst playing separately and together. Our results revealed that, for both infant and adult, and across both play conditions, motion was present over 95% of the time in experimental sessions. This therefore represents continuous contamination of the EEG signal collected during such periods. For both adult and infant, the most commonly occurring types of motion were small hand and large arm movements, which is unsurprising given that the paradigm involved object-oriented play. When further considering the types of motions that occurred as a function of play condition (Joint or Solo), we found that the majority of the adult’s most common motions (including neck, lip, jaw, arm movements and postural changes) occurred more frequently during Joint Play than Solo Play, especially large arm motions. However, the picture was reversed for infants where an increase in motion frequency was observed (e.g. chewing and nodding movements) during Solo Play relative to Joint Play. This pattern suggests that adults made more gestures during Joint Play in order to engage their child in play (de Barbaro et al., 2013). Conversely, infants who were actively engaged in play with their parents were less distracted and therefore showed a lower tendency to move.

## 3 STUDY 2

Study 2 aimed to document the EEG signal distortion generated by the commonly occurring infant and adult motions observed in Study 1. Accordingly, we repeatedly elicited selected motions from Study 1 from an adult and an infant actor whilst their EEG was recorded. This motion-contaminated EEG was then compared to their resting state EEG. For the adult, we selected the following artifacts to be modelled: **Facial Movements (FMs)**, comprising **Lip** and **Jaw** movements; **Limb Movements (LMs)**, comprising **Hand** and **Arm** movements; and finally, **Postural Movements (PMs)**, comprising **Up Neck** movements, **Nodding** (neck going down); **Leaning Forward** and **Leaning Back**. Although Lip, Neck Up and Nodding movements did not differ between the two play conditions observed in Study 1, they occurred over 10% of the time in the JP condition and therefore their effect on the adult’s EEG signal was of interest. For the infant, we selected Facial movements that were observed in Study 1 to differ between play conditions including **Talking/Babbling** and **Chewing** movements. **Limb Movements (LMs)** for the infant included **Hand**, **Arm**, **Foot** and **Leg**. These were again selected due to their high frequency of occurrence. Finally, **Postural Movements (PMs)** for the infant were **Up Neck** movements and **Nodding**.

### 3.1 Methods

#### 3.1.1 Participants

One female adult (aged 35 years) and her male infant (18 months) participated in the study. The adult was a native English speaker. She reported no neurological problems and normal hearing and vision for herself and for her infant.

#### 3.1.2 Materials

The same objects and toys were used as in Study 1. The videos from Study 1 were used as a reference to help the adult participant to imitate and reproduce the motions as accurately as possible.

#### 3.1.3 Protocol

Modelling was performed by an adult and infant to generate multiple isolated and stereotypical exemplars of each motion. The elicitation methods for Facial, Limb and Postural motions are summarised in Tables IIa and IIb for the adult and infant respectively (see Supplementary Materials Sections 3.1 and 3.2 for full descriptions). All motions were modelled by the adult participant with between 20 to 56 repetitions (Table IIa) and were recorded sequentially. Each adult motion lasted between 0.5 and 3-4 seconds. In contrast, the infant’s motions were elicited as he was naturally interacting with objects or watching a video. The infant’s motions had a variable number of repetitions, ranging from 24 to 193 iterations (Table IIb). However, the infant’s motions varied more in length than the adults’, lasting from just 0.25 seconds to tens of seconds (e.g. when the same motion, such as leg kicking, was continuously repeated). For motion artifact analysis, only periods with obvious head or body motions were selected for analysis. Insufficient Postural movements were elicited from the infant, and so no results are presented for this category of motion.

EEG during movement was compared with EEG recording during a baseline (resting state (RS)) condition. These RS measurements were acquired from the adult in a separate continuous recording of 450 seconds during which the participant was instructed to relax with her eyes open and fixated on a cross at the centre of a laptop screen (Lenovo ThinkPad, 13”, 1440 × 900 resolution, 15° angle), and to avoid any intentional movement. She was not instructed to avoid blinking or other reflex motions. Her feet, arms and shoulders were rested on cushions. For the infant, resting state measurements were obtained during periods of quiet relaxation, such as when sitting quietly in a high chair whilst watching a video. The total length of concatenated resting state segments from the infant’s recoding was 328 seconds.

#### 3.1.4 Video and EEG acquisition

##### Video recordings

As in Study 1, a webcam was used to record the adult and infant (30 frames per second). Afterwards, each recording was manually coded to ascertain the exact start and end times of the modelled behaviours.

##### Video coding

The infant’s and adult’s motion timings were extracted from each video using an identical procedure. A video-coder noted the onset and offset time of each motion by looking through the recorded video frame by frame. Only motions lasting for longer than 250ms were included for analysis. An additional inclusion criteria was that only one motion should be present at any time – periods containing overlapping motions were excluded from the analysis. For the infant’s resting state measurement, a similar approach was taken to select only periods when motion was completely absent. For the adult’s resting state measurement, the video-coder confirmed that no overt motions were present at any time during the EEG recording.

##### Video-EEG synchronisation

Video recordings were synchronised to the EEG signal by sending triggers via a radio frequency transmitter which marked the EEG trace and produced a light signal that was visible on the video recording. Synchronisation was performed manually by recording the exact frame at which the onset of the synchronisation light signal occurred. Thus, the synchronisation accuracy was limited to the temporal resolution of the video frame rate, which was 30 frames per second (33 ms).

##### EEG acquisition

A 32-channel BIOPAC Mobita mobile amplifier was used with an Easycap electrode system for both the infant and adult. Data were acquired using Acqknowledge 5.0 software, at a 500 Hz sampling rate. The ground electrode was affixed to the back of the neck as this location is the least invasive for infants.

##### EEG pre-processing

Noisy channels with raw amplitude fluctuations above 100 V above the rest of the channels for over 25% of the recording session were rejected (infant recordings: no channels rejected; adult recordings: 0-3 channels rejected across all modelled motions). Next, the data were re-referenced to the average of the remaining channels. EEG segments containing each type of motion were concatenated, creating separate continuous datasets for each motion type, and for resting state. This concatenated data was then visually inspected for eye-blinks and high amplitude fluctuations, which were removed unless directly arising from the modelled action. Finally, both motion-contaminated and resting state data were divided into 1.0s epochs for analysis.

#### 3.1.5 Analysis of motion-related EEG artifacts

For clarity of reporting, differences between a given motion-related artifact and resting state (RS) are reported for pre-defined frequency bands. The frequency bands for the adult data were Delta (1-3 Hz), Theta (3-7 Hz), Alpha (8-12 Hz), Low Beta (13-15 Hz), and High Beta (16-20 Hz). As infant oscillations are generally slower than that of their functional equivalents in adults (Orekhova et al, 1999), downward-adjusted frequency ranges were used for infants: Delta (1-3 Hz), Theta (3-6 Hz), Alpha (6-9 Hz), Low Beta (9-13 Hz), and High Beta (13-20 Hz). First, we present the average power (amplitude squared) of the EEG signal in each frequency band, averaged over all epochs, for all electrodes. Next, we assessed statistically significant differences between each artifact and resting state. For each electrode channel, and in each frequency band, an uncorrected paired t-test was conducted between the mean power of the artifact and that of the resting state signal. To ensure comparability, an equal number of 1.0s epochs was randomly selected for the resting state condition as was available for the artifact. Random epoch selection was conducted 100 times (with replacement) and only electrode sites that were significant in over 95% of permutations are reported as statistically significant.

### 3.2 Results

Examples of the raw EEG signal of artifacts produced by motion are provided in Section 3.3 (Figures S2 and S3) of the Supplementary Materials.

#### 3.2.1 Adult motion artifacts

##### 3.2.1.1 Scalp topographies of adult motion artifacts

Figure 5 shows the scalp EEG topographies for resting state power (5a) as well as for each motion artifact (5b-d), in each frequency band. During resting state, we noted that the adult showed significant central midline theta neural activity, and also high power over frontal electrodes, particularly in the high beta band (15-20 Hz), which likely corresponded to oculomotor activity. This type of oculomotor activity can be removed from the data using the techniques (e.g. ICA) that we describe in the introduction. However we chose not to apply any correction techniques to the resting state data in the present instance to avoid biasing the subsequent comparative analyses with movement artifact data. Accordingly, Figure 6 plots Cohen’s D as a measure of the t-test effect size for the difference in power between each artifact and resting state, at each electrode site, for each frequency band. Electrode sites which show a statistically significant difference in power are plotted in colour. Hotter colours (i.e. red) indicate an *increase* in power for the motion-related artifact relative to resting state.Colder colours (i.e. blue) indicate a *decrease* in power for motion-related artifact relative to resting state.

**Figure 5.**
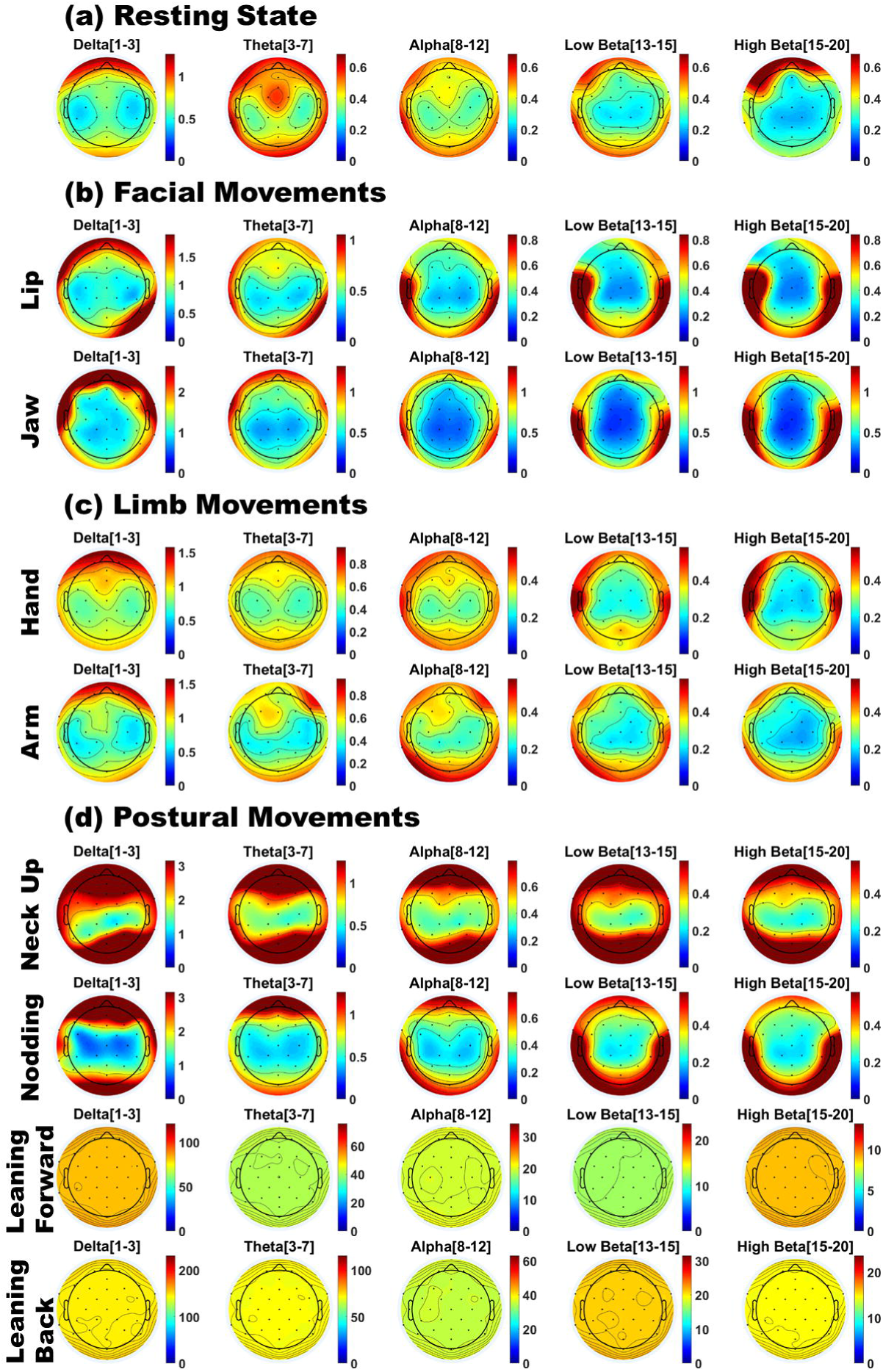
Scalp topographies of adult EEG power for (a) Resting state; (b) Facial movements; (c) Limb movements; and (d) Postural movements. Red indicates a region of high power, and blue indicates a region of low power.

**Figure 6.**
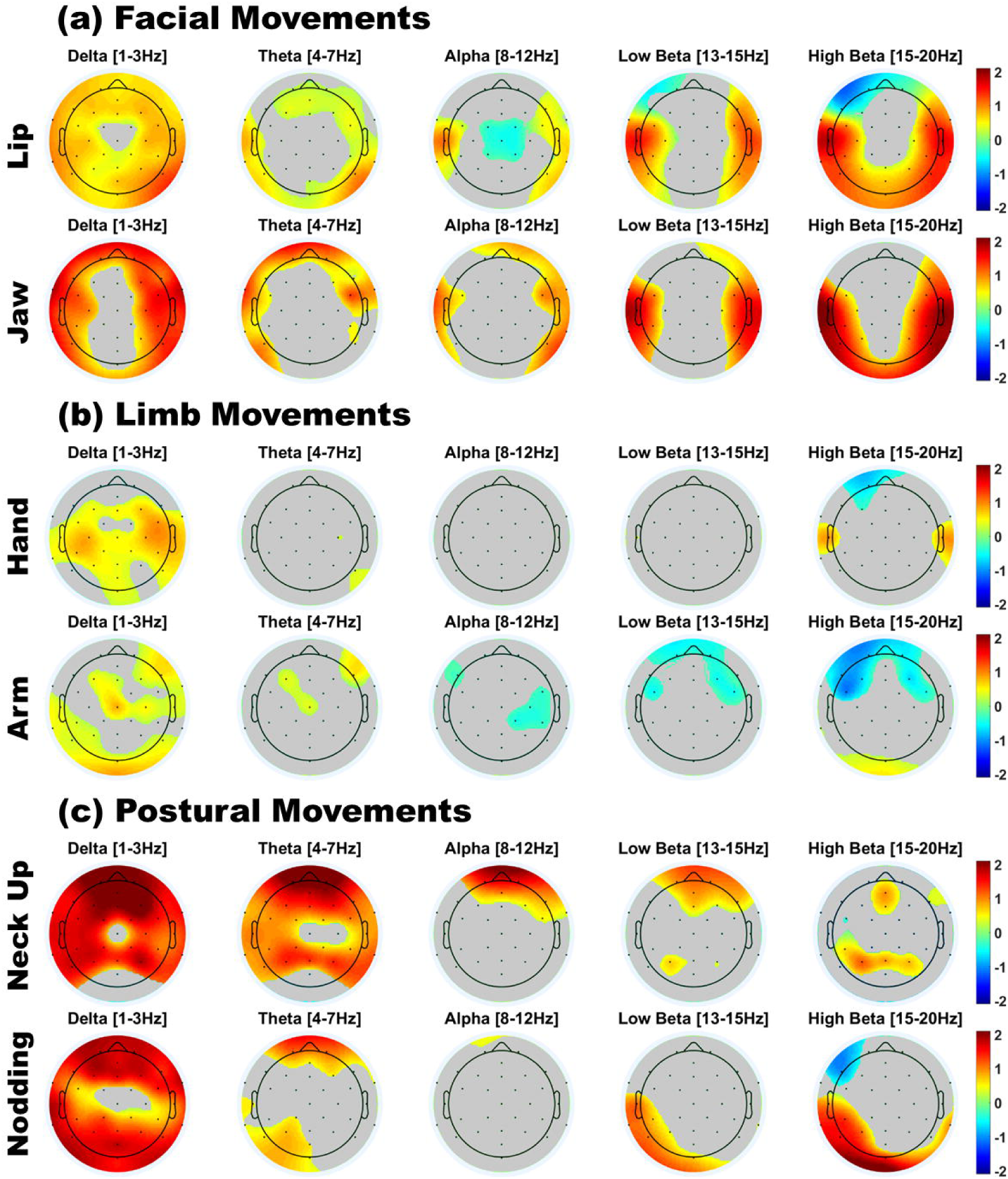
Plots of Cohen’s D-values for statistically significant differences in adult EEG power between motion artifacts and resting state (RS). In each sub-plot, only scalp regions where significant power differences were observed (p < .05) are shown in colour. Grey = non-significant result; red = higher power in the artifact; blue = higher power in RS. Leaning Forwards and Leaning Back produced very large amplitude increases (over 50 Vs across all bands), which arose because performing these motions involved synchronized displacement of the ground channel, therefore these data have not been included here.

###### Facial movements (lip, jaw)

As shown in Figure 5b and 6a, both facial movements generated large increases in power over left and right temporoparietal channels across all frequency bands, particularly in the delta and beta bands. By contrast, however, both types of facial movement produced little or no significant increase in power (as compared to resting state) over the centro-parietal region. Alpha band desynchronization (reduction in alpha power) over central electrodes was significantly higher during Lip movements as compared to resting state.

###### Limb movements (hand, arm)

As shown in Figure 5c and Figure 6b, limb movements produced relatively smaller power changes than facial movements. Compared to resting state, both hand and arm motions generated significant increases in delta power, particularly over central electrodes. Further, arm movements (and to a lesser extent, hand movements) produced a decrease in beta power over bilateral fronto-temporal electrodes, as well as some decreases in alpha power. Uniquely, hand movements elicited a bilateral increase in beta power over temporal electrodes.

###### Postural movements (neck up, nodding, leaning forward, leaning back)

All postural movements produced large power increases across all frequency bands over frontal, occipital and parieto-occipital channels. Leaning Forwards and Leaning Back, in particular, produced very large amplitude increases (over 50 Vs across all bands), which arose because performing these motions involved synchronized displacement of the ground channel. As this is a technical artifact that is specific to the current study, it will not be discussed further, and it is recommended that another site for the ground electrode should be considered for future studies that involve leaning movements. As shown in Figure 6c, both Neck Up and Nodding movements produced large increases in delta (and theta) power over almost all channels. By contrast, there were smaller and more localised increases in power observed in alpha and low beta bands, which tended to be confined to frontal or posterior (occipital) regions.

##### 3.2.1.2 Frequency spectra of adult motion artifacts

The previous sections focused on *topological* differences in the activation patterns (power) elicited by adult motion artifacts relative to resting state, in each of 5 frequency bands. Here, we present an alternative representation the same data which permits inspection of the full frequency spectrum (0-20 Hz) of each motion artifact at each electrode (rather than a band average). For example, Figure 7 shows the frequency spectra of the Lip movement artifact (black line) relative to the Resting State frequency spectra (red line) for all electrodes. The shading around the black line (artifact spectrum) represents its 95% confidence interval, and a black asterisk indicates a significant difference between the motion artifact spectrum and the resting state spectrum at that particular frequency. Similar frequency plots for each motion artifact are provided in Section 3.4 of the SM (Figure S4). Consistent with the previous topological plots, this spectral representation of the Lip movement artifact confirms that there were the largest increases in power at bilateral temporal-parietal sites (e.g. T7, T8, TP9, TP10 where power was significantly elevated for almost all frequencies). This representation also confirms that alpha power *decreased* (relative to resting state) at central sites (e.g. Cz, FC1, FC2).

**Figure 7.**
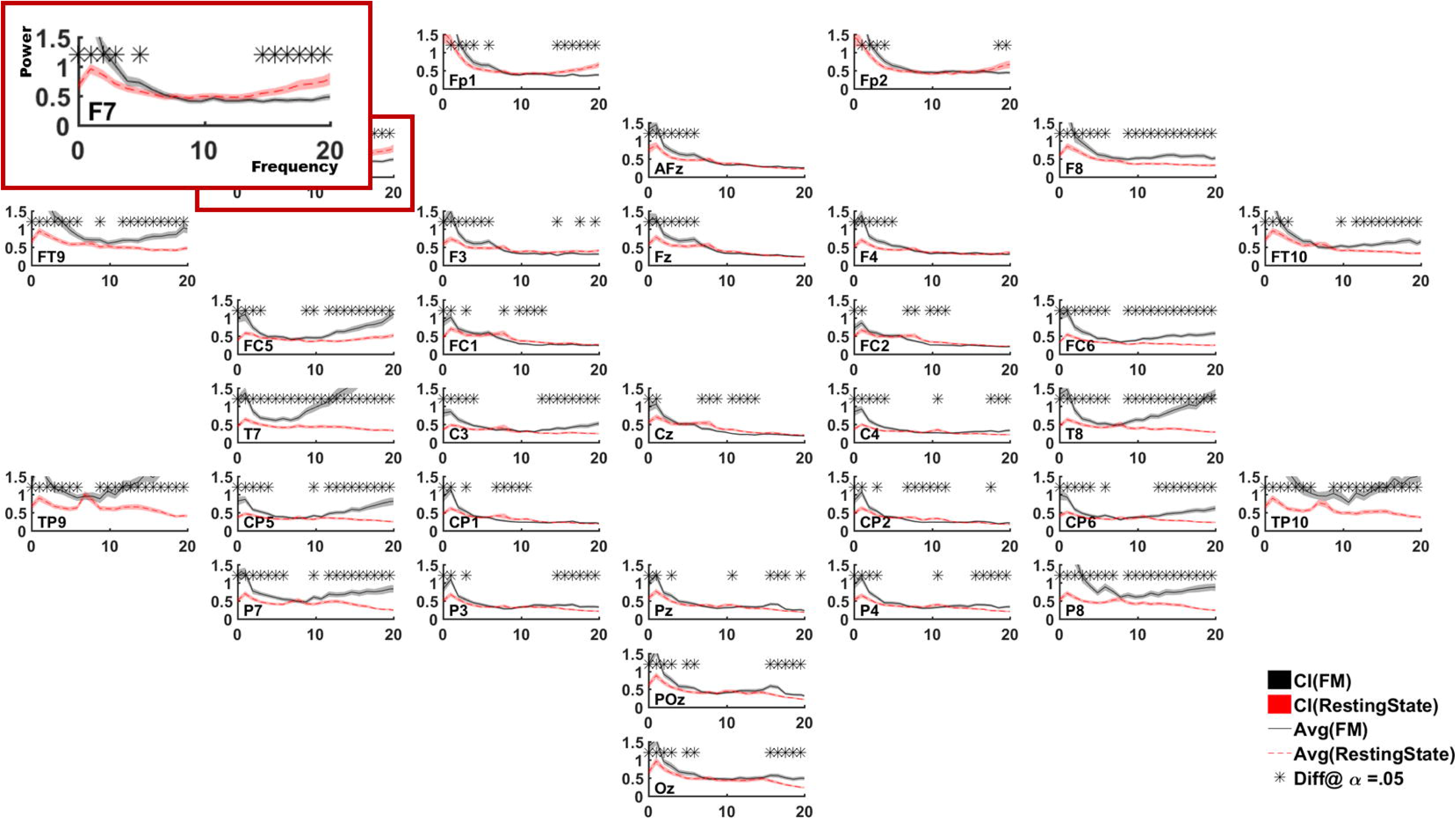
Frequency spectrum of adult facial lip movement artifact (black line) and resting state (red line) EEG for each electrode site. Shading represents 95% confidence intervals and asterisks (*) identify frequencies where a significant difference between conditions was observed. The spectra for electrode F7 (top left) is highlighted as an example.

##### 3.2.1.3 Interim summary (adult motion artifacts)

Table III provides a summary of the results across all frequency bands and channel groups. Table III is organized by frequency band and by spatial regions of interest (groups of adjacent EEG channels). “++” and “--” indicate significantly higher or lower power respectively in the motion artifact relative to resting state, observed in more than half of the electrode channels and/or frequencies within that region and band. Similarly, “+” and “-” indicate that significant differences were observed in less than half of the corresponding channels or frequencies. “O” indicates that no significant differences were observed between the motion artifact and resting state within that region and frequency band.

In general, movement artifacts were observed to generate large and significant increases in adult spectral power relative to resting state, especially in Frontal (Peripheral) and Peripheral channels, and particularly in delta and high beta frequency bands. However, there were notable exceptions in the Central and Centro-Parietal regions, as well as at POz and Oz. For these regions, we frequently noted no significant difference in power between adult motion artifacts and resting state in theta, alpha and lower beta bands. Additionally, a decrease in alpha power was sometimes observed in motion artifacts relative to resting state in central and centro-parietal channels, potentially reflecting alpha desynchronisation.

#### 3.2.2 Infant motion artifacts

##### 3.2.2.1 Scalp topographies of infant resting state and motion artifacts

###### Resting state

The topology of infant resting state EEG was characterised by high alpha power across centro-parietal regions (Figure 8a), consistent with previous reports of the infant “mu rhythm” (Cuevas et al, 2014). We also observed strong power in the low and high beta bands over bilateral temporal regions, and in the delta band over posterior and midline electrodes. As these beta and delta activation patterns were also observed across all movement measurements (see Figure 8b, 8c), we inferred that these patterns reflected tonic (and not readily observable) muscular activity in our infant participant (i.e. jaw/neck tension).

###### Facial movements (talking/babbling, chewing)

The two forms of facial movement produced highly similar scalp topographies (see Figure 8b). However, only Talking/Babbling produced a significant decrease alpha power over central, parietal and frontal regions relative to resting state (see Figure 9a).

###### Limb movements (hand, arm, foot, leg)

Upper limb (hand and arm) movements produced larger decreases in central alpha power than lower limb (foot and leg) movements (see Figures 8c and 9b), Additionally, upper limb movements increased delta power over bilateral fronto-temporal regions but this was not observed for lower limb movements. Hand and arm movements also differed from each other, with hand movements producing much stronger and more widespread decreases in alpha and beta power over central scalp regions (Figure 9b).

**Figure 8.**
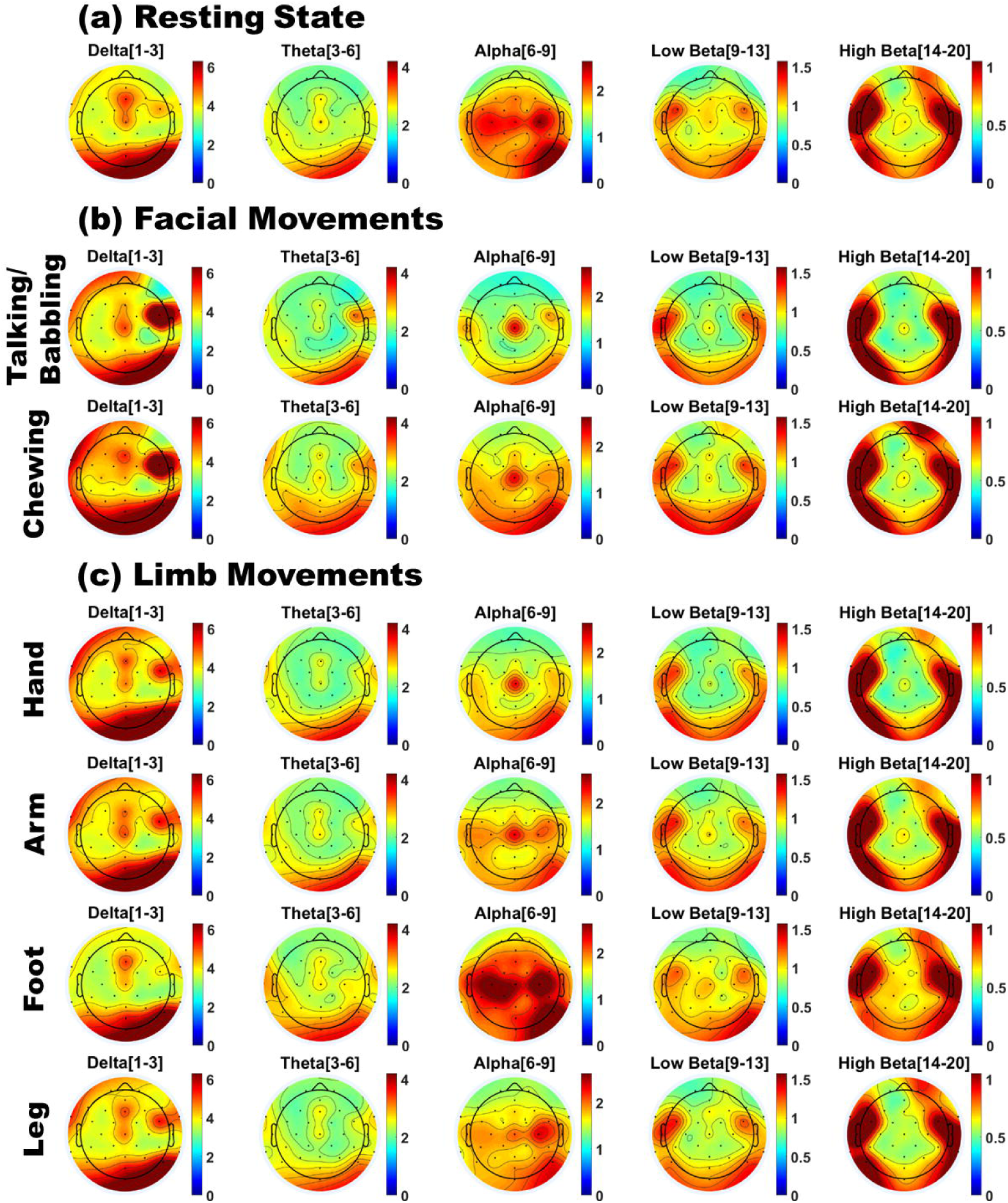
Scalp topographies of infant EEG power for (a) Resting state; (b) Facial movements; (c) Limb movements; Red indicates a region of high power, and blue indicates a region of low power.

**Figure 9.**
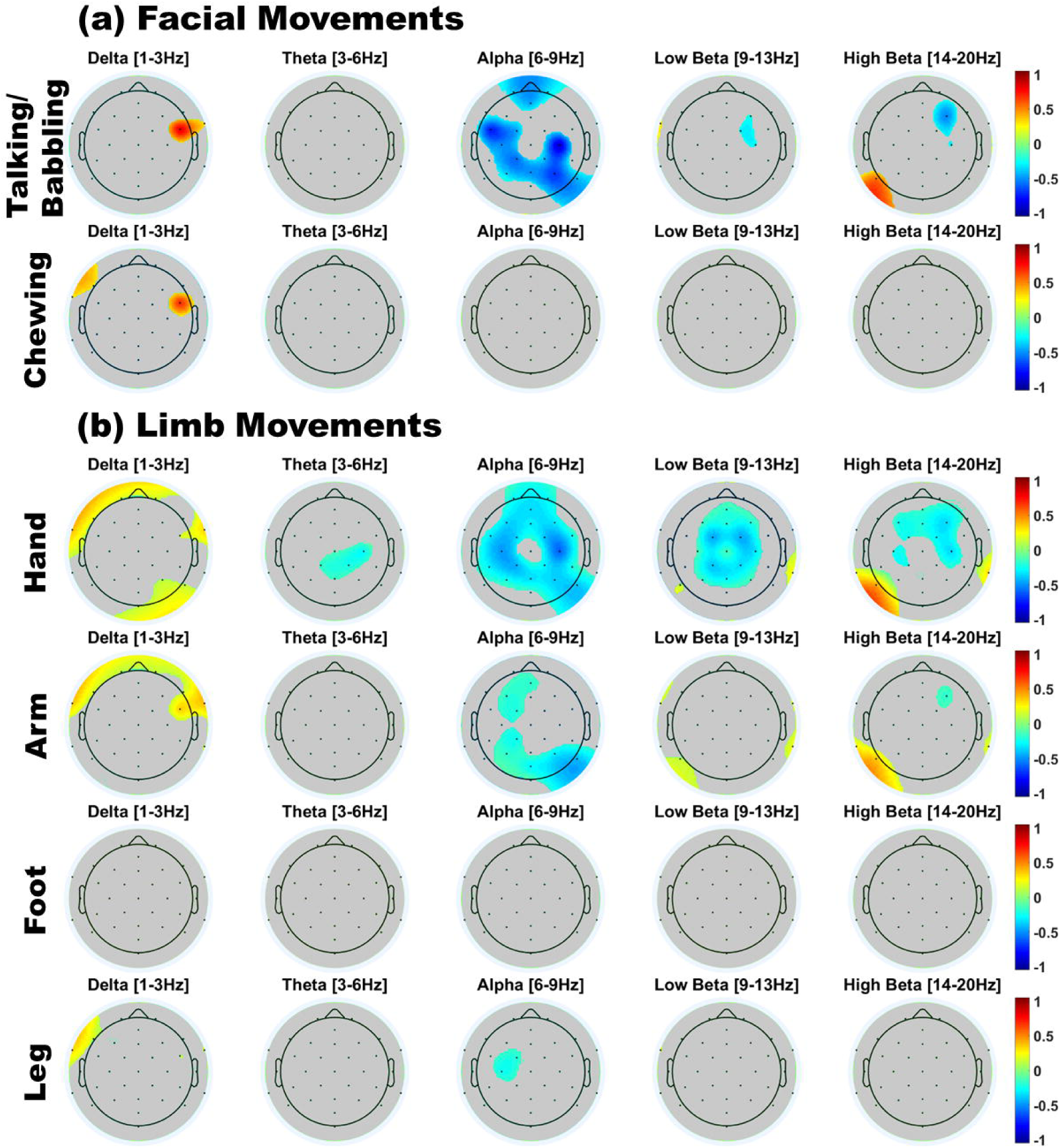
Plots of Cohen’s D-values for statistically significant differences in infant EEG power between motion artifacts and resting state. In each sub-plot, only scalp regions where significant power differences were observed (p<.05) are shown in colour. Grey = nonsignificant result; red = higher power in the artifact; blue = higher power in RS.

##### 3.2.2.2 Frequency spectra of infant motion artifacts

Figure 10 shows an illustrative spectral plot for the infant’s Talking/Babbling movements (SM Figure S5 shows equivalent plots for other movements). This clearly depicts the infant’s resting state alpha peak (red line) which was attenuated over central regions during talking/babbling (black line).

**Figure 10.**
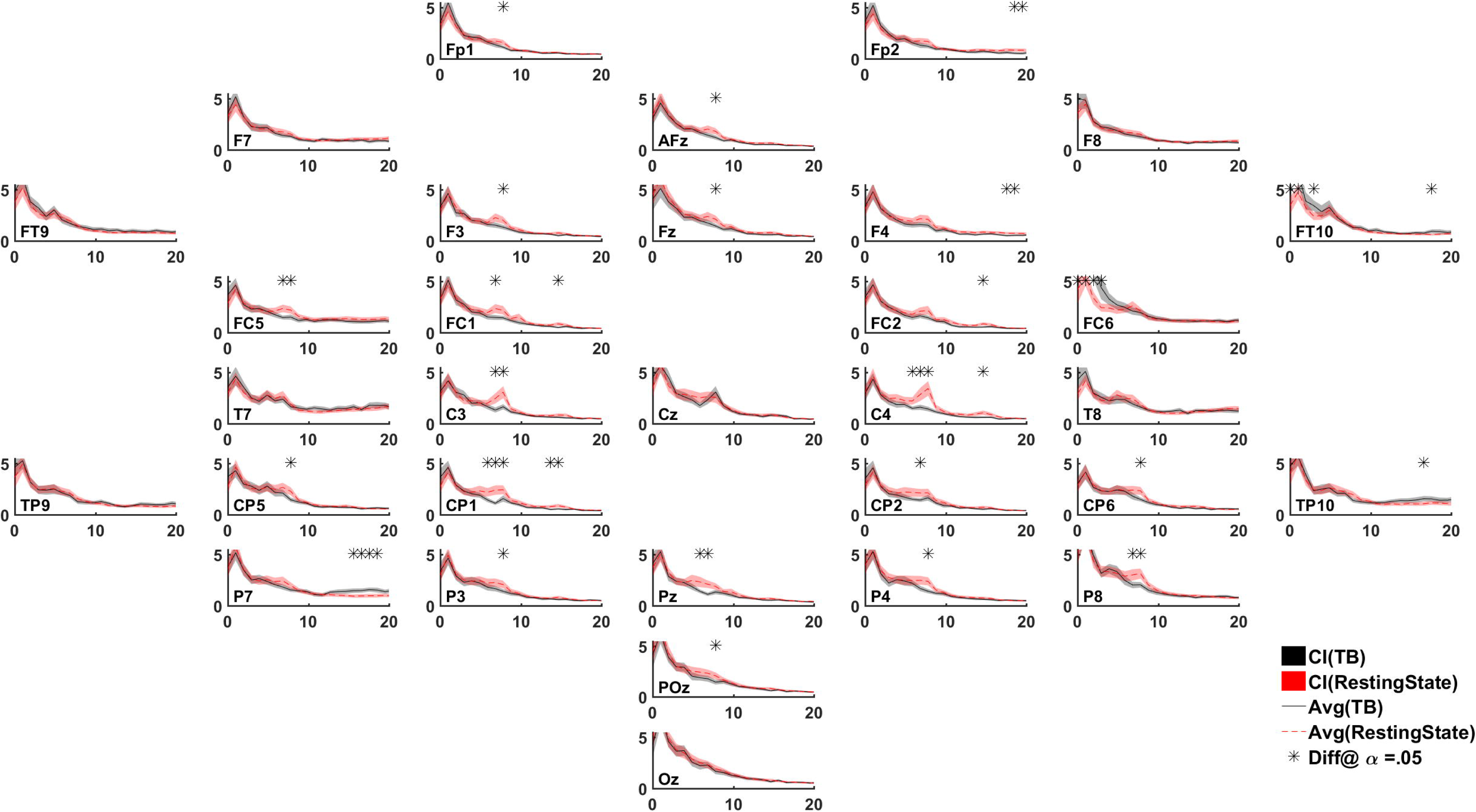
Frequency spectrum of infant talking/babbling artifact (black line) and resting state (red line) EEG for each electrode. Shading represents 95% confidence intervals and asterisks (*) identify frequencies where a significant difference between conditions was observed.

##### 3.2.3.3 Interim summary (infant motion artifacts)

Table IV shows a summary of results across all frequency bands and channel groups for the infant. Unlike the adult, the infant’s motions generated only a few significant deviations from the resting state power spectrum. Further, the significant changes related mainly to *decreases* in alpha and beta power, which were observed mainly over fronto-central, central and centro-parietal regions for Talking/Babbling and upper limb (Hand, Arm) movements. *Increases* in spectral power were more rarely observed and tended to occur in peripheral scalp regions in delta and beta bands. Finally, it is worth noting that for the infant, with the exception of peripheral channels and central channels for hand movements, the theta band did not show significant contamination by any of the modelled face and limb motions. Similarly, lower limb (foot, leg) movements produced only minimal spectral distortion with respect to the infant’s resting state activity.

## 4 STUDY 3

Study 2 examined how experimentally-induced motion artifacts affected adults’ and infants’ EEG. Here, we wished to examine whether these effects also generalised to *naturally occurring* motion patterns. To do this, we tracked and analysed infants’ spontaneous movements during a continuous EEG recording. We specifically focussed on infants for Study 3 as EEG motion artifacts in this population have not yet been systematically documented.

### 4.1 Methods

#### 4.1.1 Participants

Fourteen infants participated in the study. There were 6 boys and 8 girls in the group, with an average age of *338.85* days (SD = *59.59)*. Two infants produced insufficient resting state data due to fussiness, and so were excluded from the analysis. The remaining 12 infants comprised 5 boys and 7 girls, with an average age of *325.5* days (SD = *51.77)*. All infants were exposed to at least 50% of native English speech in their home environment. All mothers reported no neurological problems and normal hearing and vision for their infants.

#### 4.1.2 Materials

The stimuli used were a series of brief videos that were suitable for infants. The videos included familiar nursery rhymes (such as ‘Twinkle Twinkle Little Star’) that were sung by a female adult, interspersed with different (static) cartoon pictures. All infants saw the same set of videos, presented in a counterbalanced order. The videos lasted up to *22.77* minutes in total.

#### 4.1.3 Protocol

Here, infants’ spontaneous motions during passive video viewing were analysed. This differed from Study 2, where individual motions were specifically elicited and modelled. As shown in Figure 11, infants passively watched videos while seated in a high chair, with their mothers seated adjacent to them. A camera recorded infants’ behaviour (at 30 frames per second) throughout the session. Afterwards, each video was manually screened frame-by-frame and coded to ascertain the start and end times of each motion type, using the same criteria as Study 2. Based on the quantity of naturally occurring motions produced by infants, sufficient exemplars were collected for analysis of **Jaw (Chewing), Hand, Arm, Foot** and **Leg** movements.

**Figure 11.**
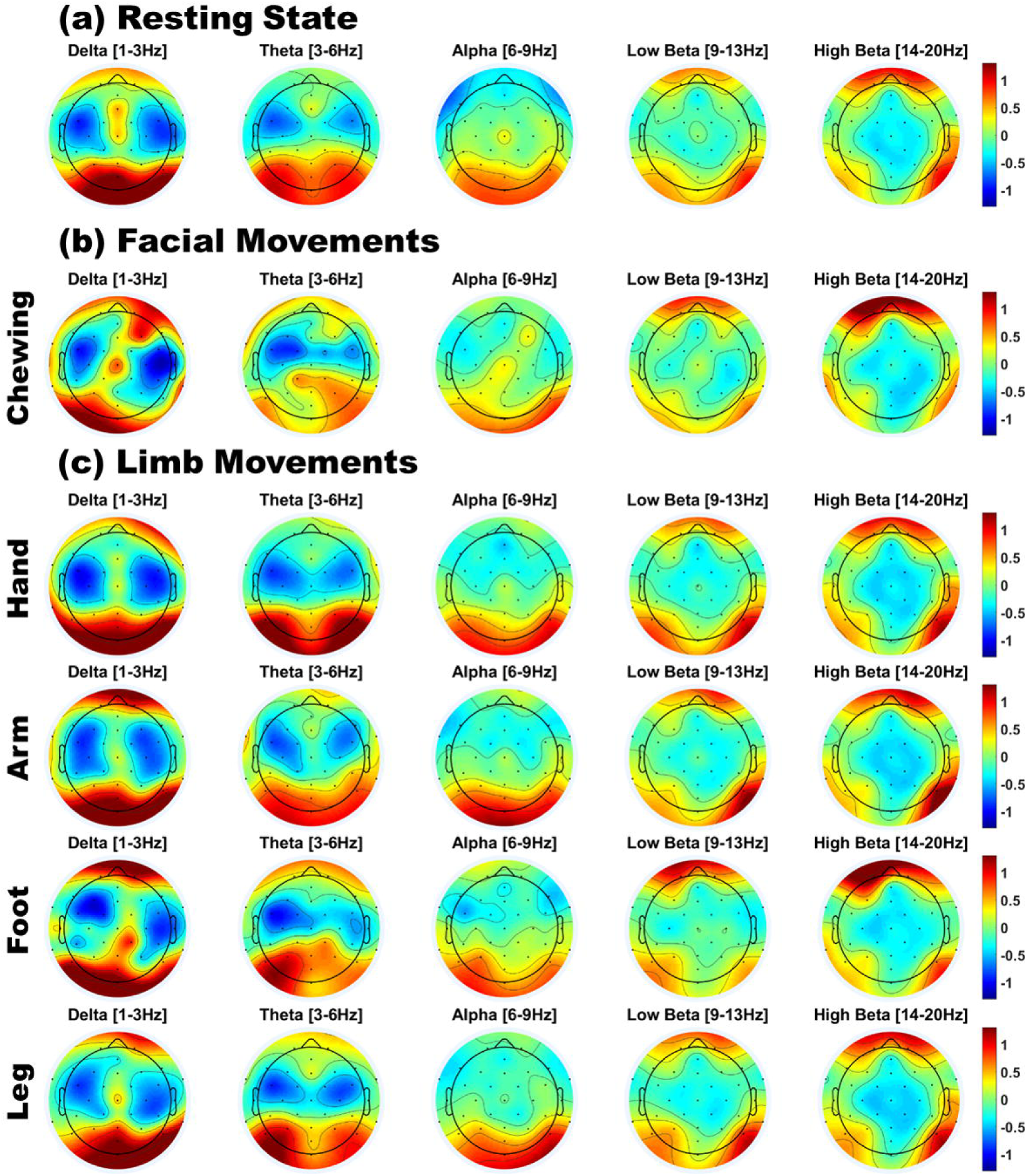
Experimental Setup for Study 3. Infants were seated on a highchair next to their mothers. A camera, placed in a central location in front of participants, recorded infants’ behaviour and motions. Panel A (left) illustrates resting state behaviour, when the infant showed no visible motion. Panel B (right) illustrates leg movement by the infant. Written consent was obtained for the use of these images.

##### Motions

Only isolated (i.e. non-overlapping) and stereotypical exemplars of each motion were used for analysis. The total duration of each type of (naturally-occurring) motion contributed by each infant is shown in Table V. All 12 infants whose data were analysed produced motion in at least two out of five motion categories. As not all the infants spontaneously produced all types of motion, the number of participants analysed for each motion type varied between 4 and 10 (see Table V).

##### Resting state

Resting state periods were strictly defined as periods during which the infant exhibited fixated gaze with no visible facial or bodily motion and maintained this state for at least half a second. Periods with any visible motion, including eye-blinks, were excluded. All infants included in the analysis produced at least *10* seconds of Resting State data (average of 38.3 seconds).

#### 4.1.4 EEG acquisition and analysis

The EEG system, acquisition protocol and pre-processing steps were identical to those used in Study 2. Table V shows the number of rejected channels for each infant, as well as the total duration of EEG (divided into 1-second epochs) used for analysis for each condition. As in Study 2, analysis was conducted using infant-appropriate EEG frequency bands: Delta (1-3 Hz), Theta (3-6 Hz), Alpha (6-9 Hz), Low Beta (9-13 Hz), and High Beta (13-20 Hz). For each EEG frequency, we computed the average Resting State power (amplitude squared) of the EEG signal, averaged over all epochs, for all electrodes, for all infants. Next, statistically significant differences in mean power (averaged across epochs) between each motion type and resting state were assessed at the group level by conducting frequency point-wise t-tests on the z-normalized mean difference (Motion - RS), for each EEG band. The full spectral results for each channel and motion type can be seen in Figure S7 (Section 4.3 in SM). P-values were adjusted to correct for the number of channels being compared using the Benjamini-Hochberg (1995) False Discovery Rate [FDR] procedure with alpha at *.05*. For comparison, the uncorrected results are also presented in the SM Figure S6 (Section 4.2).

### 4.2 Results

#### 4.2.1 Scalp topographies

##### Resting state

As shown in Figure 12a, infants’ resting state scalp topology was characterised by high power over posterior regions in delta and theta bands, and high alpha power over centro-parietal regions. These patterns were also observed in Study 2. However, unlike Study 2, the cohort of infants tested here showed higher beta power over bilateral orbitofrontal regions, and relatively less power over bilateral temporal regions, which could reflect the presence of oculomotor activity (such as microsaccades). Individual plots for each infant’s resting state scalp topologies are presented in Figure S5a (Section 4.1 in SM).

##### Movement

Compared to resting state EEG, the scalp topology of infants’ movement EEG also showed a broadly similar pattern of high delta/theta power over posterior regions and high beta power over orbitofrontal regions (see Figure 12b and 12c). However, some significant variations in topography (by movement class) were also observed. As shown in Figure 13, upper limb movements (hand and arm) produced significant *decreases* in power over central sites, accompanied by *increases* in power at peripheral sites. These effects were particularly prominent at theta, alpha and beta frequencies. By contrast, facial and lower limb movements (foot and leg) produced virtually no significant changes as compared to infants’ resting state topography. Individual plots for each infant’s motion-related scalp topologies are presented separately for each motion type in Figures S5b through to S5f (Section 4.1 in SM).

**Figure 12.**
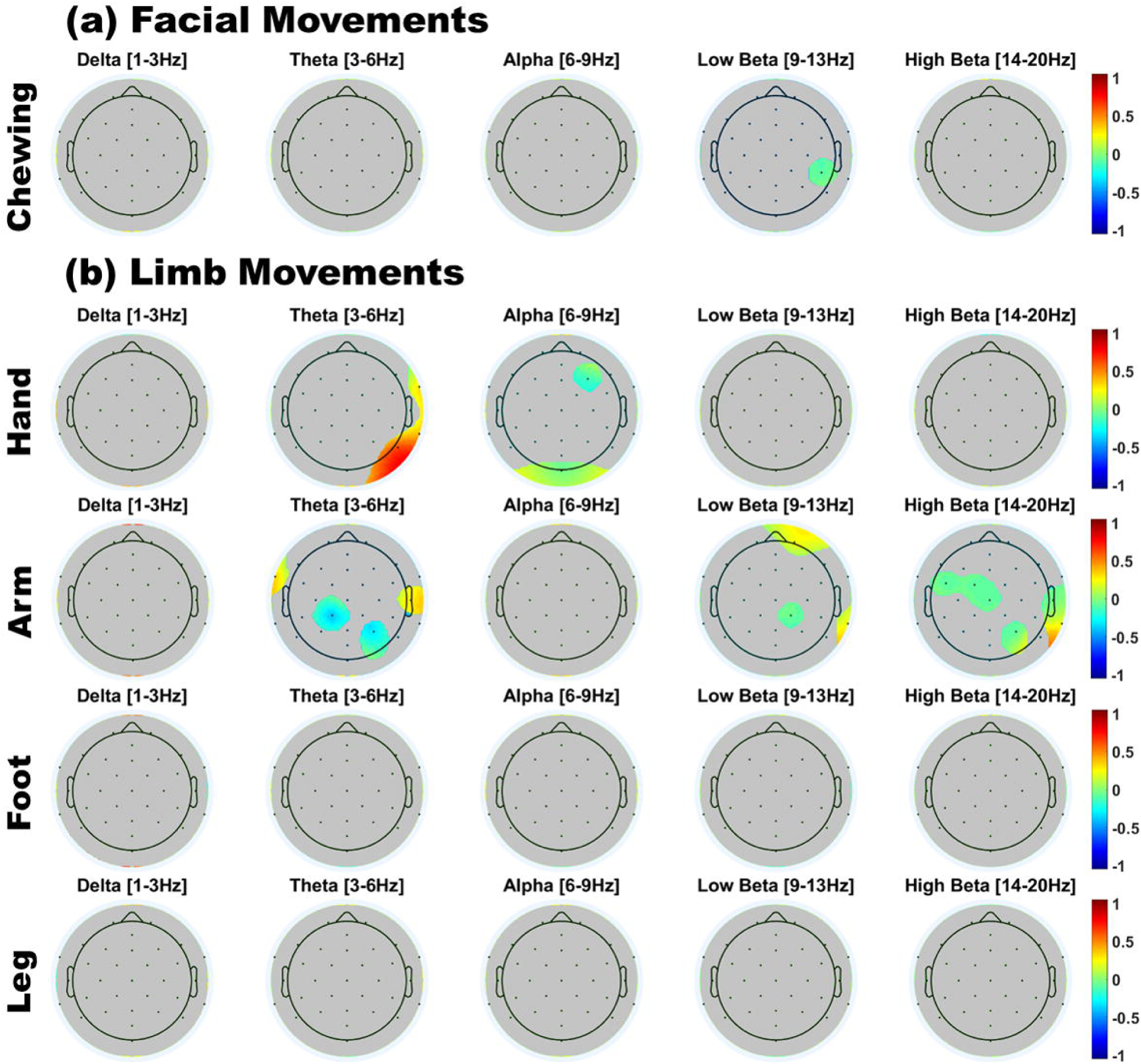
Scalp topographies of infant EEG power for (a) Resting state; (b) Facial movements; and (c) Limb movements; Power is z-normalize dpower [uV^2^/Hz], averaged over infants. Red indicates a region of above-average power, and blue indicates a region of below-average power.

**Figure 13.**
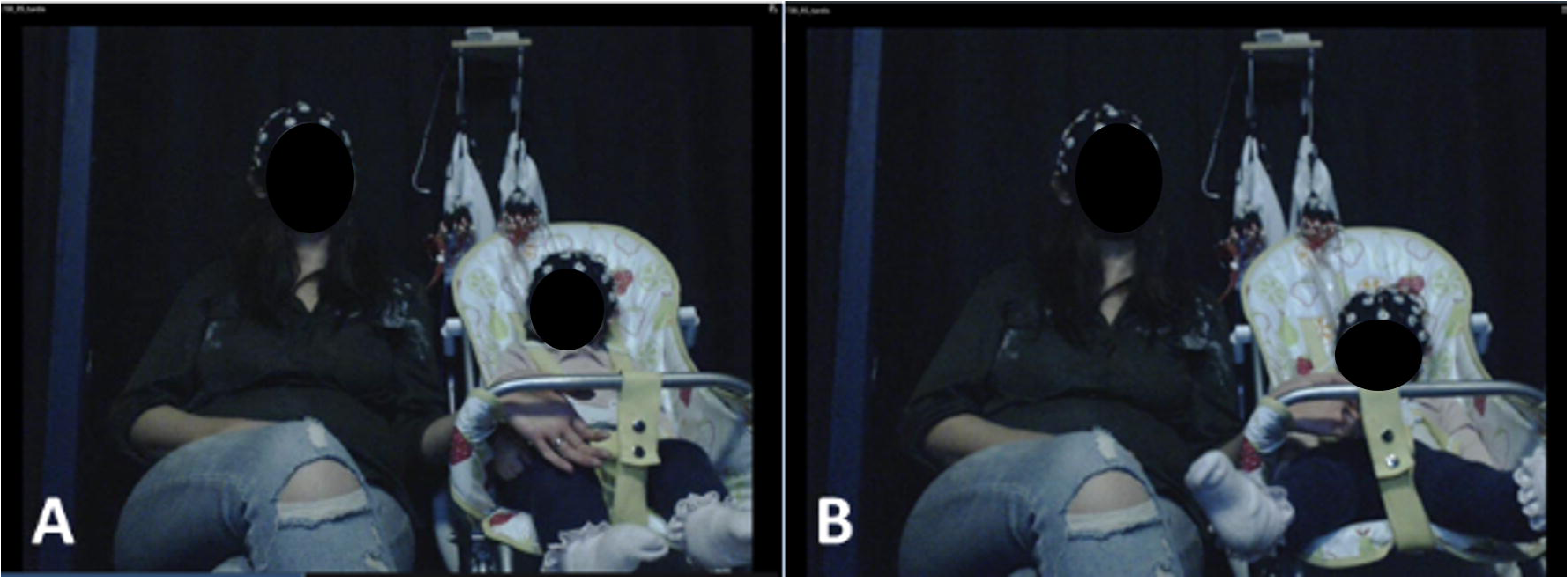
Scalp topography of significant differences (Cohen’s D-values) between each movement class and resting state. In each sub-plot, only scalp regions where significant power differences were observed (Benjamin-Hochberg FDR corrected at p<.05) are shown in colour. Grey = nonsignificant result; red = higher power during movement; blue = higher power during resting state.

#### 4.2.2 Interim summary

Consistent with Study 2, upper limb movements produced the largest effects as compared to chewing or lower limb movements. Also consistent with Study 2, upper limb movement caused decreased central theta and beta power and increased beta power at peripheral sites. However, unlike Study 2, we observed no significant changes in delta power for any motion type, and less evidence of central alpha desynchronization for hand and arm movements.

## 5 DISCUSSION

EEG recordings are highly prone to distortion by motion-related artifacts, which can result in the misinterpretation of underlying neural processes, or even the inaccurate detection and diagnosis of brain disorders (Guerrero-Mosquera et al., 2012). For example, fMRI studies have demonstrated that head movements (which are more common in young children and clinical conditions such as autism spectrum disorder) can produce biases in computations of brain functional connectivity (Deen & Pelphrey, 2012), making it difficult to dissociate true experimental/clinical differences in brain functional connectivity from artifactual effects arising from increased head motion in the scanner. Young infants present a particular challenge as they have a high natural tendency for movement, which cannot be constrained by instruction. This work represents the first systematic assessment of the effects of naturalistic (social) human motion on the recorded EEG signal – contrasting adult and infant motion artifacts. It is intended that this work will build toward a more comprehensive database or ‘Artifact Library’ which could later serve as a common resource for EEG researchers in social and developmental neuroscience.

Study 1 assessed the prevalence of motion in adult-infant dyads during social and non-social naturalistic play paradigms. We observed that motion occurred during >95% of the time, for both infants and adults, and in both types of social play settings. Interestingly, whilst adults’ gestural motions (e.g. arm movements) tended to increase in frequency for social as compared to non-social play, infants showed less looking around and chewing. This may be attributable to the fact that infants were more engaged and less distracted during social play with their mothers. For such datasets, it would not be feasible to adopt a simple approach of rejecting (excluding) all motion-contaminated data, as this would entail losing an unacceptably high proportion of data. However, before artifact removal methods (such as ICA or CCA) can be effectively applied to the EEG signal, it is first necessary to understand the exact distortion that these motions would produce. Accordingly, Study 2 and 3 attempted to document the topographical and spectral properties of each of the most prevalent (frequently-occurring) types of motion that were observed in Study 1. In Study 2, this was achieved with the assistance of adult and an infant “actors”, who provided repeated exemplars of facial, limb and postural motion whilst their EEG was recorded. In Study 3, spontaneously-occurring motion by infants was analysed. The spectral properties of these motion-contaminated signals were then contrasted against a resting state baseline.

Study 2 showed that for the adult, *all* types of movement (facial, limb and postural) generated significant changes in spectral power relative to resting state. Across all types of motion, the strongest effects occurred in peripheral scalp regions (including frontal and temporal sites), particularly in delta and high-beta frequency bands. A few exceptions were noted in central and centro-parietal channels (e.g. C3 and C4), as well as at POz and Oz. For these regions, motion produced no (or few) significant changes in power in theta, alpha and low-beta frequency bands. In particular, we replicated results from previous studies on adult speech production artifacts (e.g. Brooker and Donald, 1980; Ganushchak and Schiller, 2008; Laganaro and Perret, 2011). Muscle artifact contamination was observed to be greatest over frontal and temporal scalp regions and to be less severe over central regions, particularly at frequencies over 12 Hz. In our study, we also observed significant jaw artifact contamination in the delta band (1-3 Hz), suggesting that the common practice of low-pass filtering under 12 Hz to remove speech production artifacts may be inadequate.

By contrast, the infant’s motions generated only a few significant deviations from the resting state power spectrum. These changes related mainly to *decreases* in alpha and beta power (alpha desynchronization) over central scalp regions. More rarely, *increases* in spectral power tended to occur in peripheral regions, and were observed in delta and beta bands. The results of Study 3 (naturally-occurring motion from N=12 infants) were consistent with the infants results from Study 2 (elicited motion, N=1) in two main ways. First, upper limb movement (hand and arm) produced stronger and more widespread artifacts than chewing and lower limb movements (foot and leg). Second, upper limb movement artifacts were characterised by a *reduction* in power at central sites, accompanied by *increases* in power at peripheral sites. However, unlike Study 2, these changes occurred primarily at theta and beta frequencies, with less prominent/no effects in the alpha and delta bands (see also Appendix Figure S6 for results with uncorrected p-values). Accordingly, we conclude that the most generalisable and pernicious effects of infant motion on the EEG signal are observed in relation to upper limb movement. Further, in general, infant movement artifacts appear to be most likely to cause reductions in central power at theta, alpha and beta frequencies in conjunction with increases in delta and beta power at peripheral electrodes.

One potential explanation for the apparent lower motion contamination in infants’ as compared to adults’ EEG data could be that insufficient data was collected from infants to reveal true differences. However, this is unlikely as, in Study 2, the total amount of resting state data was comparable for infant (328s) and adult (450s); and the infant also had a higher number of recorded instances for all motion types. An alternative explanation could be that inter-trial variability was higher in infant than adult motions. However, in Study 2 there was no noticeable difference either in the variability of raw data (upon visual inspection), nor in the 95% confidence intervals around the FFT mean for every channel and across all frequencies, as can be seen in Section 3.4 of the Supplementary Materials. Further, Study 3, which was conducted with a larger number of infants, confirmed the results from Study 2 – namely that the effects of motion on the infant EEG signal were relatively limited. However, it should be noted that infants’ resting state data differed qualitatively from the adult’s resting state data. Since we could not instruct infants to produce a state of rest, their resting state data was collected incidentally (e.g. whilst watching a video) and periods of non-motion were identified through video coding. This protocol of recording continuous EEG and selecting relevant segments offline is frequently used in infant research (e.g. Orekhova et al., 2014). However, it is possible that this procedure inadvertently included some tonic muscle activity that was not visible on video, leading to an underestimation of the true extent of the effects of motion on the infant’s EEG signal.

Alpha or “mu rhythm” suppression has been well-documented in infants and young children in relation to both action production and action observation (Liao et al, 2015). For example, Marshall et al (2011) reported suppression effects in the 6-9 Hz range, broadly distributed over the scalp, when 14-month old infants were engaged in action observation in a social context. This raises the question of whether the movement-related suppression effects observed here are truly “artifactual”, since they could also reflect the cognitive or social processes that underpin infants’ action generation. According to this view, only unintentional or involuntary movement produces truly artifactual effects. However, to definitively separate these effects would require concurrent measures of intentionality and cognitive processing, along with active and passive manipulations of participant motion, which are beyond the scope of the current study.

### 4.1 Implications for EEG research using naturalistic paradigms

The growth of naturalistic EEG paradigms reflects the view that movement is a natural neural state (Makeig et al, 2009) and that cognitive processes themselves are embodied (Gramann et al, 2011). The brain maintains representations of its internal (proprioception) and external (motor behaviours and audio-visual scene) environment – and these representations are constantly, dynamically updated through action. To study cognition in this holistic and action-oriented way, new technologies and imaging methods are required, such as mobile sensors that can synchronously image the brain (i.e. wireless EEG) and the body (i.e. motion capture) and perform online registration of the two modalities. One such system is the sensor technology for mobile brain/body imaging (MoBI) which has shown promising results for studying changes in the EEG power spectrum in relation to participants’ gait during locomotion (Gramann et al, 2011). The MoBI system has also been used to localise independent components in relation to motions such as head-turning, pointing, and walking in a 3D virtual reality orientation task (Makeig et al, 2009). However, when such sophisticated methods for tracking and removing EEG artifacts are not available, precautions should be taken when analysing data from naturalistic paradigms, as discussed next.

#### Adult EEG

Given the large and pernicious effects of all classes of motion on the adult EEG signal, it is important that any data collected during naturalistic social paradigms is interpreted with great care. It may be advisable to completely exclude the most heavily affected electrode sites from analyses (e.g. peripheral frontal and temporal channels), rather than to attempt artifact removal. Depending on the type of motion that occurred during the paradigm, the signal from the remaining central, parietal and/or occipital channels may further benefit from selective bandpass filtering (e.g. to exclude delta and high-beta frequencies), before more targeted artifact removal methods are applied.

#### Infant EEG

Our results indicated that for infants, one major effect of jaw and limb (particularly upper limb) motion was to reduce EEG power in the theta and alpha bands. Accordingly, if EEG researchers are investigating phenomena where infant alpha band effects are predicted (e.g. mu de-synchronisation or frontal alpha asymmetry), care must be taken to avoid including confounding jaw or limb motions, which can independently create changes in alpha band power across frontal and central sites. For example, jaw motions may provide a confounding factor when infants suck on a teething toy whilst watching screen-based experimental stimuli. It is important that experimenters check, and report (e.g. by video coding) the relative occurrence of these movement artifacts in infant participants across experimental conditions and groups, to verify that any reported results cannot be explained by differences in patterns of movement. Further, like the adult data, we also found that infants’ peripheral EEG channels were particularly vulnerable to motion-related power increases, and therefore recommend particular caution (or exclusion) when analysing these channels.

### 4.2 Limitations and Future Directions

The major limitation to the current work is that the studies were conducted with a small sample size (N=10 for Study 1, N=2 for Study 2, N=12 for Study 3). This limits the wider generalisability of these findings, as individuals may differ substantially in their motion patterns, and also in the effects of motion on their EEG signals. Nonetheless, the current work is an initial, but necessary, first step toward a better understanding of the effects of motion on adult and infant EEG data during naturalistic paradigms. Further studies with a larger number of participants, and a wider range of modelled motions will be necessary to ascertain the extent to which these effects are generalisable, and to inform the future development of methods for EEG artifact removal.

## ACKNOWLEDGEMENTS

This research was funded by a UK Economic and Social Research Council (ESRC) Transforming Social Sciences Grant ES/N006461/1 (to V.L. and S.W.), a Nanyang Technological University start-up Grant M4081585.SS0 (to V.L.), an ESRC Future Research Leaders Fellowship ES/N017560/1 (to S.W.), and a Rosetrees Medical Trust PhD Studentship A1414 (to S.G.).All the authors declare no conflicts of interest.

## BIBLIOGRAPHY

Babiloni, F., Astolfi, L., Cincotti, F., Mattia, D., Tocci, A., Tarantino, A., et al. (2007). Cortical activity and connectivity of human brain during the prisoner’s dilemma: an EEG hyperscanning study. In Engineering in Medicine and Biology Society, 2007. EMBS 2007. 29th Annual International Conference of the IEEE (pp. 4953–4956). IEEE. 10.1109/IEMBS. doi: 2007.4353452

Batty, M., Meaux, E., Wittemeyer, K., Rogé, B., Taylor, M. J. (2011). Early processing of emotional faces in children with autism: an event-related potential study. Journal of experimental child psychology, 109: 4. doi.org/10.1016/j.jecp.2011.02.001

Bell, M. A., Cuevas, K. (2012). Using EEG to Study Cognitive Development: Issues and Practices. Journal of Cognition and Development, 13: 3. doi.org/10.1080/15248372.2012.691143

Benjamini, Y. and Hochberg, Y. (1995). Controlling the false discovery rate: A practical and powerful approach to multiple testing. Journal of the Royal Statistical Society, Series B, 57, 289–300.

Brooker, B. H., Donald, M. W. (1980). Contribution of the speech musculature to apparent human EEG asymmetries prior to vocalization. Brain and Language, 9: 2. doi.org/10.1016/0093-934X(80)90143-1

Chaumon, M., Bishop, D. V., Busch, N. A. (2015). A practical guide to the selection of independent components of the electroencephalogram for artifact correction. Journal of neuroscience methods, 250. doi.org/10.1016/j.jneumeth.2015.02.025

Cohen, M. X. (2014). Analyzing neural time series data: theory and practice. MIT Press. 51–4

Cuevas, K., Cannon, E. N., Yoo, K., Fox, N. A. (2014). The Infant EEG Mu Rhythm: Methodological Considerations and Best Practices. Developmental Review, 34: 1. doi.org/10.1016/j.dr.2013.12.001

Dahl, A. (2017). Ecological Commitments: Why Developmental Science Needs Naturalistic Methods. Child Development Perspectives, 11: 2. doi: 10.1111/cdep.12217

De Barbaro, Kaya, Christine M. Johnson, and Gedeon O. Deák (2013). Twelve-month ‘‘social revolution' 'emerges from mother-infant sensorimotor coordination: A longitudinal investigation, Human Development, 56(4), 223–248.

Deen, B., Pelphrey, K. (2012). Perspective: brain scans need a rethink. Nature, 491(7422), S20–S20. doi:10.1038/491S20a

De Haan, M. (Ed.). (2013). Infant EEG and event-related potentials. Psychology Press.

Dumas, G., Lachat, F., Martinerie, J., Nadel, J., George, N. (2011). From social behaviour to brain synchronization: review and perspectives in hyperscanning. Irbm, 32: 1. doi.org/10.1016/j.irbm.2011.01.002

Dumas, G., Nadel, J., Soussignan, R., Martinerie, J., Garnero, L. (2010). Inter-brain synchronization during social interaction. PloS one, 5: 8. doi:10.1371/journal.pone.0012166

Dumas, G., Laroche, J., Lehmann, A. (2014). Your body, my body, our coupling moves our bodies. Frontiers in human neuroscience, 8. doi.org/10.3389/fnhum.2014.01004

Fabes, R. A., Martin, C. L., Hanish, L. D., Updegraff, K. A. (2000). Criteria for Evaluating the Significance of Developmental Research in the Twenty-First Century: Force and Counterforce. Child Development, 71: 1. doi: 10.1111/1467-8624.00136

Ganushchak, L.Y., Schiller, N.O. (2008). Motivation and semantic context affect brain errormonitoring activity: an event-related brain potentials study. NeuroImage, 39. doi.org/10.1016/j.neuroimage.2007.09.001

Ganushchak, L., Christoffels, I., Schiller, N. O. (2011). The use of electroencephalography in language production research: a review. Frontiers in psychology, 2. doi.org/10.3389/fpsyg.2011.00208

Gramann, K., Gwin, J. T., Ferris, D. P., Oie, K., Jung, T. P., Lin, C. T., … Makeig, S. (2011). Cognition in action: Imaging brain/body dynamics in mobile humans. Reviews in the Neurosciences, 22(6), 593–608. https://doi.org/10.1515/RNS.2011.047

Guerrero-Mosquera, C., Navia-Vazquez, A., Trigueros, A. M. (2012). EEG signal processing for epilepsy. INTECH Open Access Publisher

Gwin, J. T., Gramann, K., Makeig, S., Ferris, D. P. (2010). Removal of movement artifact from high-density EEG recorded during walking and running. Journal of Neurophysiology, 103: 6. doi:10.1152/jn.00105.2010

Islam, M. K., Rastegarnia, A., Yang, Z. (2016). Methods for artifact detection and removal from scalp EEG: a review. Neurophysiologie Clinique/Clinical Neurophysiology, 46: 4. doi.org/10.1016/j.neucli.2016.07.002

Klados, M. A., Papadelis, C., Braun, C., Bamidis, P. D. (2011). REG-ICA: a hybrid methodology combining blind source separation and regression techniques for the rejection of ocular artifacts. Biomedical Signal Processing and Control, 6: 3. doi.org/10.1016/j.bspc.2011.02.001

Kuhl, P. K. (2010). Brain mechanisms in early language acquisition. Neuron, 67: 5. doi.org/10.1016/j.neuron.2010.08.038

Laganaro, M., Perret, C. (2011). Comparing electrophysiological correlates of word production in immediate and delayed naming through the analysis of word age of acquisition effects. Brain Topography, 24: 1. doi.org/10.1007/s10548-010-0162-x

Lawhern, V., Hairston, W. D., McDowell, K., Westerfield, M., Robbins, K. (2012). Detection and classification of subject-generated artifacts in EEG signals using autoregressive models. Journal of neuroscience methods, 208: 2. doi:10.1016/j.jneumeth.2012.05.017. ISSN 0165-0270

Liao, Y., Acar, Z. A., Makeig, S., & Deak, G. (2015). EEG imaging of toddlers during dyadic turn-taking: Mu-rhythm modulation while producing or observing social actions. NeuroImage, 112, 52–60. https://doi.org/10.1016/j.neuroimage.2015.02.055

Lindenberger, U., Li, S. C., Gruber, W., Müller, V. (2009). Brains swinging in concert: cortical phase synchronization while playing guitar. BMC neuroscience, 10: 1. doi.org/10.1186/1471-2202-10-22

Maguire, M. J., Magnon, G., Fitzhugh, A. E. (2014). Improving data retention in EEG research with children using child-centered eye tracking. Journal of Neuroscience Methods, 238. doi.org/10.1016/j.jneumeth.2014.09.014

Makeig, S., Gramann, K., Jung, T. P., Sejnowski, T. J., & Poizner, H. (2009). Linking brain, mind and behavior. International Journal of Psychophysiology, 73(2), 95–100. https://doi.org/10.1016/j.ijpsycho.2008.11.008

Marshall, P. J., Young, T., & Meltzoff, A. N. (2011). Neural correlates of action observation and execution in 14-month-old infants: An event-related EEG desynchronization study. Developmental Science, 14(3), 474–480. https://doi.org/10.1111/j.1467-7687.2010.00991.x

McMenamin, B. W., Shackman, A. J., Maxwell, J. S., Bachhuber, D. R., Koppenhaver, A. M., Greischar, L. L., Davidson, R. J. (2010). Validation of ICA-based myogenic artifact correction for scalp and source-localized EEG. Neuroimage, 49: 3. doi.org/10.1016/j.neuroimage.2009.10.010

Molla, M. K. I., Islam, M. R., Tanaka, T., Rutkowski, T. M. (2012). Artifact suppression from EEG signals using data adaptive time domain filtering. Neurocomputing, 97. doi.org/10.1016/j.neucom.2012.05.009

Nathan, K., Contreras-Vidal, J. L. (2015). Negligible motion artifacts in scalp electroencephalography (EEG) during treadmill walking. Frontiers in Human Neuroscience, 9. doi.org/10.3389/fnhum.2015.00708

Noris, B., Nadel, J., Barker, M., Hadjikhani, N., Billard, A. (2012). Investigating gaze of children with ASD in naturalistic settings. PloS one, 7: 9. doi.org/10.1371/journal.pone.0044144

O'Regan, S., Faul, S., and Marnane, W. (2010). Automatic detection of EEG artefacts arising from head movements. In Engineering in Medicine and Biology Society (EMBC), 2010 Annual International Conference of the IEEE (pp. 6353–6356). IEEE. doi:10.1109/IEMBS.2010.5627282

Orekhova, E. V, Elsabbagh, M., Jones, E. J., Dawson, G., Charman, T., & Johnson, M. H. (2014). EEG hyper-connectivity in high-risk infants is associated with later autism. Journal of Neurodevelopmental Disorders. https://doi.org/10.1186/1866-1955-6-40

Plöchl, M., Ossandón, J. P., König, P. (2012). Combining EEG and eye tracking: identification, characterization, and correction of eye movement artifacts in electroencephalographic data. Frontiers in human neuroscience, 6. doi.org/10.3389/fnhum.2012.00278

Porcaro, C., Medaglia, M. T., Krott, A. (2015). Removing speech artifacts from electroencephalographic recordings during overt picture naming. NeuroImage, 105. doi.org/10.1016/j.neuroimage.2014.10.049

Reis, P. M., Hebenstreit, F., Gabsteiger, F., von Tscharner, V., Lochmann, M. (2014). Methodological aspects of EEG and body dynamics measurements during motion. Towards a New Cognitive Neuroscience: Modeling Natural Brain Dynamics, 9. doi.org/10.3389/fnhum.2014.00156

Reynolds, G. D. (2015). Infant visual attention and object recognition. Behavioural Brain Research, 285, 34–43.

Richards, J. E., Reynolds, G. D., Courage, M. L. (2010). The Neural Bases of Infant Attention. Current Directions in Psychological Science, 19: 1

Saby, J. N., Marshall, P. J. (2012). The Utility of EEG Band Power Analysis in the Study of Infancy and Early Childhood. Developmental Neuropsychology, 37: 3. doi.org/10.1080/87565641.2011.614663

Shackman, A. J., McMenamin, B. W., Slagter, H. A., Maxwell, J. S., Greischar, L. L., Davidson, R. J. (2009). Electromyogenic artifacts and electroencephalographic inferences. Brain topography, 22: 1. doi.org/10.1007/s10548-009-0079-4

Smith, M. L. (1982). Benefits of naturalistic methods in research in science education. Journal of Research in Science Teaching, 19: 8. doi: 10.1002/tea.3660190802

Stroganova, T. A., Orekhova, E. V., Posikera, I. N. (1999). EEG alpha rhythm in infants. Clinical Neurophysiology, 110: 6. doi.org/10.1016/S1388-2457(98)00009-1

Sweeney, K. T., Ward, T. E., McLoone, S. F. (2012). Artifact removal in physiological signals—Practices and possibilities. IEEE transactions on information technology in biomedicine, 16:3. doi: 10.1109/TITB.2012.2188536

Tamis-LeMonda, C. S., Kuchirko, Y., Luo, R., Escobar, K., Bornstein, M. H. (2017). Power in methods: Language to infants in structured and naturalistic contexts. Developmental Science. doi: 10.1111/desc.12456

Telkemeyer, S., Rossi, S., Nierhaus, T., Steinbrink, J., Obrig, H., Wartenburger, I. (2011). Acoustic processing of temporally modulated sounds in infants: evidence from a combined near-infrared spectroscopy and EEG study. Frontiers in psychology, 2. doi: 10.3389/fpsyg.2011.00062

Teplan, M. (2002). Fundamentals of EEG measurement. Measurement science review, 2: 2).

Tunnell, G. B. (1977). Three dimensions of naturalness: An expanded definition of field research. Psychological Bulletin, 84: 3. doi: 10.1037/0033-2909.84.3.426

Turnip, A., Junaidi, E. (2014, August). Removal artifacts from EEG signal using independent component analysis and principal component analysis. In Technology, Informatics, Management, Engineering, and Environment (TIME-E), 2014 2nd International Conference on (pp. 296–302). IEEE. doi: 10.1109/TIME-E.2014.7011635

Yuval-Greenberg, S., Tomer, O., Keren, A. S., Nelken, I., Deouell, L. Y. (2008). Transient induced gamma-band response in EEG as a manifestation of miniature saccades. Neuron, 58:3. doi.org/10.1016/j.neuron.2008.03.027

Widrow, B., & Stearns, S. D. (1985). Adaptive signal processing (Vol. 15). Englewood Cliffs, NJ: Prentice-Hall

